# Robustness to nuisance perturbations enables unsupervised evaluation of single-cell foundation models

**DOI:** 10.64898/2026.08.22.746357

**Authors:** Ahmed Sallam, Jesse Gillis

**Affiliations:** University of Toronto

## Abstract

Single-cell foundation model evaluations have relied almost exclusively on downstream tasks. While these tasks measure whether an embedding recovers annotated cell types, batches, or trajectories, they cannot determine if that structure is reproducible or merely an artifact of a single noisy draw, a key limitation since incomplete sampling is intrinsic to single-cell measurement. Here, we introduce a fully unsupervised evaluation framework grounded in a fundamental principle: a faithful representation must preserve its neighbourhood structure under nuisance perturbations that mimic technical and sampling variation. Across five scFMs, a PCA baseline, and 39 datasets, we show that models ranked as near-equivalent by standard benchmarks differ nearly twofold in local neighbourhood preservation under a perturbation discarding just 5% of counts. This structural instability is scale-dependent and often masked by visually coherent embeddings. Cluster-level stability under resampling tracks established bio-conservation metrics (Spearman ρ=0.78), showing that invariance to nuisance perturbations captures representation quality no benchmark measures directly.

## Main

Single-cell foundation models (scFMs) are trained on millions of cells to learn general-purpose representations of cell state, transferable to new datasets with little or no additional training^1–4^. These models encode each cell as an embedding, a low-dimensional vector positioned so that cells in similar states lie close together. In practice, scFMs are used primarily for these embeddings, which serve as input to algorithms for downstream analysis including clustering, perturbation prediction, and trajectory inference^5–8^. The premise is that a single pretrained representation can be shared across these analyses, rather than fitting a method to each dataset.

Whether scFMs deliver on this premise remains contested, largely due to the absence of a gold standard. In natural language processing, although evaluation is expensive, human readers can serve as a direct judge, and benchmarks approximate that judgment at scale^9^. Single-cell transcriptomics instead adopted quantitative measurement because no one can characterize a count matrix by inspection, leaving no expert whose judgment a benchmark can stand in for. Current evaluations therefore measure performance on downstream tasks such as cell type annotation, batch integration, and perturbation prediction, which require labels to define a ground truth^10–14^. Those labels are themselves typically produced through a pipeline in which cells are clustered and then annotated^15^, so evaluating an embedding against them asks, in part, whether it recovers a partition produced by a previous analysis. Task performance additionally conflates representation quality with task difficulty, metric choice, and annotation scheme, and model rankings can shift with the datasets, metrics, and baselines chosen^10,14,16–19^. Independent evaluations have accordingly reached differing conclusions, with scFMs variously matching, exceeding, or falling short of simple baselines such as highly variable gene (HVG) selection or principal component analysis (PCA)^18–20^, and an embedding can rank first on every standard metric while distorting biological structure^21^.

Where no external gold standard exists, the properties that remain available are intrinsic ones, and the most basic is that a result should survive resampling of the data that produced it. This is a familiar principle in statistical inference, whether evaluating a statistic against a null, an estimate across bootstrap redraws, or a model across cross-validation partitions. It is particularly relevant in single-cell transcriptomics because incomplete sampling is intrinsic to the measurement^22^. A cell’s library is typically designed to be an incomplete draw from its transcriptome, and the analytical conventions of the field exist to recover signal that survives that draw. A representation that reorganizes under resampling of counts is therefore not reporting cell state but reporting a particular draw, independently of any annotation. This same principle is central to representation learning, where representations are required to remain invariant to nuisance variation while remaining selective for semantic variation^23–25^. Applying it to single-cell transcriptomics requires defining what semantic information an embedding must preserve. Because cell state is defined primarily by a cell’s relationship to surrounding cells rather than by its absolute position, and algorithms for clustering, integration, and trajectory inference all operate on nearest-neighbour relationships^5–7^, neighbourhood structure is the natural object. We therefore ask whether an embedding preserves its neighbourhood structure under nuisance perturbations that mimic technical and sampling variation. Since these perturbations are defined independently of cell-type labels or other annotations, the evaluation is unsupervised.

Recent work has begun to probe scFM representations beyond standard task benchmarks. One line introduces label-based metrics that compare an embedding’s neighbourhood graph to a reference graph derived from raw counts or principal components, flagging embeddings that score well on conventional measures yet distort biological structure^21^. Such reference graphs are themselves dataset-dependent and biased toward the structure of the original data, and they still presuppose a fixed notion of correct neighbourhoods. Closest to the present work, a recent benchmark pairs zero-shot utility with a notion of robustness by perturbing dataset structural properties such as cell number, gene number, and class balance and measuring the resulting change in downstream task performance^26^. However, robustness there is still defined at the task level, asking whether a model’s benchmark score is stable when the composition of the dataset changes, rather than whether the representation itself is invariant to nuisance variation.

Here we introduce a label-free evaluation framework grounded in a single principle: a faithful representation should preserve neighbourhood structure under nuisance perturbations that mimic technical and sampling variation (Fig. 1a). We operationalize this through controlled perturbations of the count matrix, including Poisson resampling, downsampling, gene dropout, and local smoothing. Because each manipulation maps every perturbed cell to exactly one reference cell, distortion is measured against a known correspondence rather than an external annotation. We quantify neighbourhood structure invariance at complementary scales, spanning pairwise distances, local and global neighbour rankings, and cluster assignment, and show that the resulting scores correlate with label-based measures of biological conservation, indicating that invariance to nuisance perturbations tracks biological conservation despite requiring no external annotations. Applying this framework to five scFMs and a PCA baseline across 39 datasets, we find that neighbourhood preservation varies substantially between methods, that distortion can be scale-dependent, with a representation preserving coarse global organization while disrupting local neighbourhoods, and that a representation’s response to specific perturbations aligns with its training objectives. Together, these results support evaluating scFMs through the invariance of neighbourhood structure under nuisance perturbations, a representation-level property that standard task benchmarks cannot measure directly. All raw metrics are explorable through an online dashboard (https://scfm-controlled-manipulations.streamlit.app/), with a Python implementation of perturbations and metrics at https://github.com/ahmedh-sallam/scfm-controlled-manipulations.

**Figure 1.**
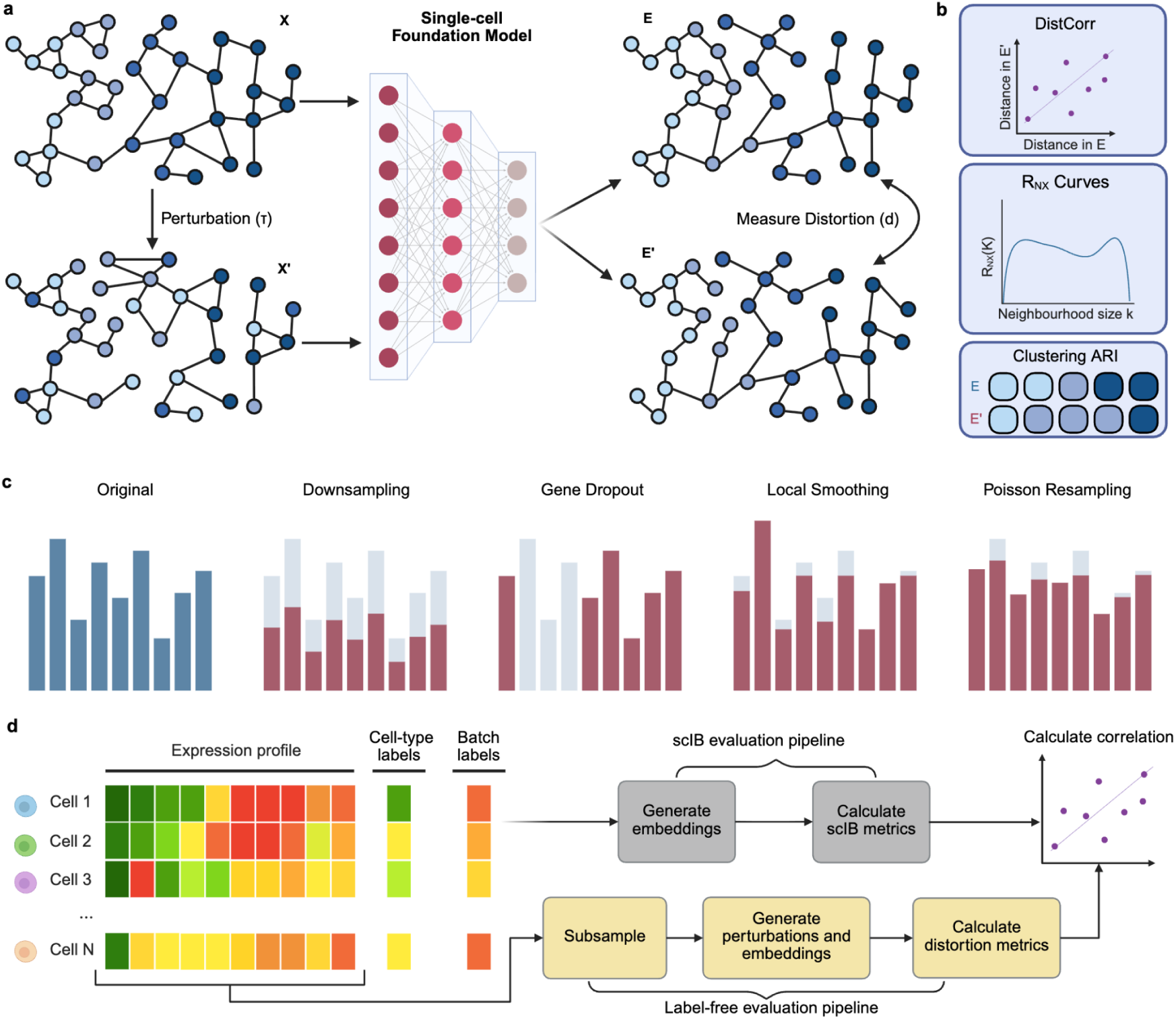
Evaluation framework. **(a)** Overview. A reference expression matrix X is embedded by a single-cell foundation model to give the reference embedding E. A perturbation τ is applied to X to give a perturbed matrix X′, embedded by the same model to give E′. Cells in X′ correspond one-to-one to cells in X, so distortion d is measured directly between E and E′ without labels. **(b)** Metrics. Distortion between E and E′ is quantified by distance correlation (DistCorr), the correlation between the pairwise distances of the two embeddings; R_NX_ curves, the rank-based agreement between neighbour rankings as a function of neighbourhood size K, from which a local score weighting small neighbourhoods and a global score weighting large neighbourhoods are derived; and clustering agreement (ARI), the adjusted Rand Index between Leiden partitions of E and E′. **(c)** Perturbations. Four nuisance perturbations, each shown as its effect on a cell’s expression profile relative to the reference. Downsampling reduces the total counts per cell, gene dropout sets a random subset of expressed genes to zero, Poisson resampling redraws each count about its observed value, and local smoothing replaces each profile with the average of its nearest neighbours. **(d)** Pipeline. A single-cell dataset comprises an expression matrix together with cell-type and batch label columns if available. The label-free branch uses expression alone, subsampling the dataset, generating perturbations and embeddings, and computing distortion metrics between the reference and perturbed embeddings. The label-based branch embeds the reference dataset and computes scIB metrics from the cell-type and batch labels. The two are correlated to establish whether stability tracks performance on established benchmarks.

### Evaluation framework for measuring neighbourhood distortion under nuisance perturbations

Methods relying on k-nearest-neighbour (kNN) graphs are ubiquitous in single-cell biology, with applications in visualization, clustering, cell-type annotation, batch correction, and trajectory inference^5–7,27,28^. These methods remain effective despite the high dimensionality of expression data, which would otherwise concentrate pairwise distances^29^, because gene co-regulation confines cells to a lower-dimensional manifold where local proximity remains meaningful^7,30–32^. We therefore quantify distortion in terms of neighbourhood relationships rather than absolute position in embedding space. Because reference and perturbed data are compared within the same embedding space, arbitrary geometric properties such as coordinate system, scale, and anisotropy are intrinsically controlled, and the resulting metrics are directly comparable across models and datasets.

The framework compares two embeddings of the same set of cells (Fig. 1a). We embed an unperturbed expression matrix (the reference), apply a perturbation to the same matrix, embed the perturbed version, and measure the degree to which its neighbourhood structure is distorted relative to the reference. Because each perturbed cell corresponds to exactly one reference cell by construction, distortion is measured against a known correspondence rather than an external annotation. We refer to the applied perturbations as nuisance perturbations, as each reflects technical or sampling variation (Fig. 1c). We consider four such perturbations: Downsampling reduces the total counts per cell to reflect variation in sequencing depth. Gene dropout sets a random subset of expressed genes to zero, mimicking incomplete transcript capture. Poisson resampling redraws each count from a Poisson distribution centred on its observed value, capturing shot noise in the counting process^22^. Local smoothing replaces each cell’s profile with the average of its nearest neighbours, mirroring operations shared by denoising and metacell methods^33,34^. Each perturbation is applied across a range of strengths to measure distortion as a continuous function of perturbation severity (Table S1).

We quantify the distortion between the reference and perturbed embeddings using four complementary metrics (Fig. 1b). Distance correlation (DistCorr) computes the correlation of pairwise distances between the two embeddings^35^. From their co-ranking matrix we derive two rank-based scores, following the R_NX_ formulation of Lee and Verleysen^36,37^ as implemented in ViScore^38^. For each cell, this calculates a normalized score analogous to a Jaccard index of shared nearest neighbours. The local score weights small neighbourhoods and measures whether each cell’s immediate neighbours are preserved; the global score weights large neighbourhoods and measures whether the coarse arrangement of cells is preserved. Clustering agreement (ARI) compares Leiden partitions of the two embeddings using the adjusted Rand Index^39^, measuring whether the discrete groupings are stable.

Invariance to nuisance perturbations alone is not sufficient for a faithful representation. Because each model is evaluated relative to its own reference embedding rather than an external ground truth, a representation completely independent of its input would be trivially invariant, lacking the selectivity required to capture genuine biological variation. We therefore compared our label-free metrics with three widely used label-based evaluations: scIB bio-conservation, scIB batch correction, and trajectory inference (Fig. 1d).

Because repeatedly perturbing and embedding large datasets is computationally expensive, we include a subsampling step for every dataset larger than 5,000 cells. We verified that this approximation was stable across six human atlases ranging from hundreds of thousands to millions of cells (Table S2). Across seven subsample sizes ranging from 200 to 20,000 cells and 18 perturbation settings, the median coefficient of variation across random subsamples fell below 0.05 for every sample size above 1,000 cells, and most perturbation–metric combinations converged by 5,000 cells (Fig. S1). We therefore use 5,000-cell subsamples throughout, giving stable invariance estimates while keeping the benchmark computationally tractable.

### scFMs differ in the degree to which they distort neighbourhood structure

Invariance to nuisance perturbations differed substantially across embedding methods. We evaluated five scFMs (scGPT, Geneformer, scFoundation, SCimilarity, scConcept^2–4,41,42^) and a PCA baseline across 39 datasets (Data S1). DistCorr was corrected against a permutation null, whereas the remaining metrics are inherently adjusted for chance (Figs. S2-S7). As expected, all four metrics decreased monotonically with increasing perturbation strength for every method (Fig. 2a). Local smoothing, however, produced a qualitatively different response from the three count-based perturbations. Under count-based perturbations, neighbourhood preservation declined approximately in proportion to perturbation strength; under downsampling, for example, the mean Local R_NX_ across models fell from 0.72 at 95% count retention to 0.43 at 40%. Local smoothing instead reached its minimum by *k* = 10 and changed by less than 0.05 as *k* increased to 100. Metric agreement also diverged under smoothing: DistCorr remained high (0.69-0.92) for all methods, while Local R_NX_ decreased substantially (0.25-0.46), because averaging cells preserves the coarse geometry captured by DistCorr while disrupting the local neighbourhoods that rank-based metrics evaluate.

**Figure 2.**
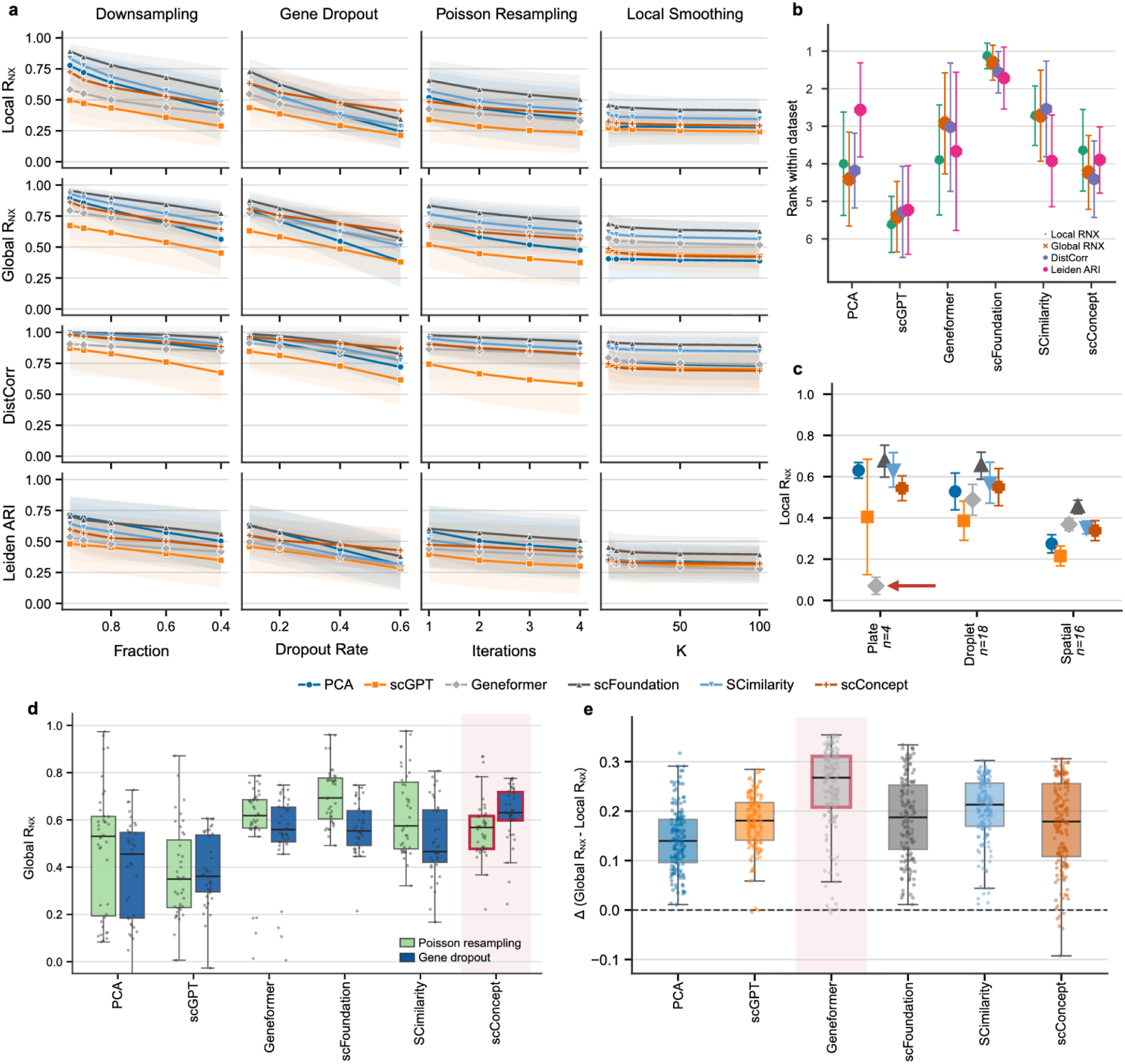
Representations differ in neighbourhood structure invariance under nuisance perturbations. Six representations were evaluated across n=39 datasets: five single-cell foundation models and a PCA baseline refit on each perturbed matrix. All values are null-corrected where applicable, and higher values indicate greater preservation throughout. **(a)** Neighbourhood structure invariance against perturbation strength, with rows corresponding to metrics and columns to perturbations. Each line represents one model averaged across datasets, shaded regions denote standard deviation (SD), and strength increases from left to right within each column. **(b)** Within-dataset ranks. For each dataset and metric, representations were ranked by perturbation-averaged preservation (rank 1 = highest preservation). Points indicate the mean rank per representation and metric, with error bars representing ±1 SD across datasets. **(c)** Local R_NX_ by assay technology, averaged over perturbations. Points represent the mean per representation and technology group, with error bars denoting ±1 SD. The arrow marks Geneformer on plate-based data. **(d)** Global R_NX_ under Poisson resampling (iterations=4) and gene dropout (fraction=0.6), with one point per dataset. scConcept is highlighted as the model preserving the most structure under gene dropout. The center line corresponds to the median, with boxes indicating the interquartile range (IQR) and whiskers extending to 1.5× IQR. **(e)** Difference between Global R_NX_ and Local R_NX_, with one point per dataset and perturbation setting. The center line corresponds to the median, with boxes indicating the interquartile range (IQR) and whiskers extending to 1.5× IQR. Geneformer is highlighted as the representation with the largest gap.

Across datasets, the degree of neighbourhood invariance differed markedly between models (Fig. 2a). scFoundation was consistently the most invariant, with perturbation curves lying above those of every other method across most metrics and perturbations. SCimilarity ranked second, and scGPT was consistently the least invariant; Geneformer, scConcept, and PCA occupied intermediate positions. These differences were substantial even under mild perturbations. For example, under downsampling to 95% of original counts, Local R_NX_ ranged from 0.89 for scFoundation and 0.84 for SCimilarity to 0.49 for scGPT, with PCA (0.78), scConcept (0.73), and Geneformer (0.58) between these extremes. Ranking models within each dataset produced the same ordering (Fig. 2b), scFoundation first (mean rank 1.1-1.7), and scGPT last (5.2-5.6) across all four metrics, both with little variation across datasets, while the remaining methods showed overlapping rank distributions that precluded a strict ordering. PCA showed the strongest metric dependence, ranking near fourth for Local R_NX_, Global R_NX_, and DistCorr but improving to 2.6 for Leiden ARI. Together, these results show that neighbourhood structure invariance consistently distinguishes the most and least invariant representations.

Neighbourhood structure invariance also depended on assay technology (Fig. 2c). All methods preserved less neighbourhood structure on spatial datasets than on plate- or droplet-based datasets (Local R_NX_ 0.21-0.45 versus 0.38-0.68). Geneformer departed from this pattern: on plate-based datasets, its Local R_NX_ fell close to zero, whereas every other method retained values between 0.40 and 0.68, including the refitted PCA baseline at 0.63. Because only four plate-based datasets were available, we regard this observation as preliminary. Nevertheless, the same assay-specific pattern was observed across all four evaluation metrics (Fig. S8). One possible explanation is Geneformer’s preprocessing pipeline, which normalizes each gene’s expression relative to the median expression in its pretraining corpus. Because this corpus is filtered to include droplet-based datasets only, the model may generalize poorly to the differing count distributions typical of plate-based assays.

Training objectives also shaped model responses to specific perturbations. scConcept was trained using a contrastive objective in which each cell is divided into two complementary views containing disjoint sets of genes, an augmentation that closely resembles our gene dropout perturbation. Consistent with this, scConcept showed the highest global neighbourhood structure invariance under gene dropout, both in absolute performance (Fig. 2d, median Global R_NX_ 0.63 versus 0.36-0.56 for the other models) and in relative change (+11% compared with Poisson resampling, versus −20% to +3% for the remaining models). The same pattern was observed for DistCorr but not for Local R_NX_ or Leiden ARI (Fig. S9). Although this observation does not establish a general relationship between training objectives and invariance, it shows that nuisance perturbations can recover representation properties that reflect how a model was trained.

### Geneformer preserves global structure while distorting local neighbourhoods

Neighbourhood structure invariance depended strongly on the scale at which it was evaluated. Across nearly every dataset and embedding method, Global R_NX_ exceeded Local R_NX_ (Fig. 2e), indicating that coarse organization is more invariant to nuisance perturbations than immediate neighbourhood relationships. Geneformer showed the strongest scale dependence, with a larger gap between Global and Local R_NX_ than any other method (median 0.27 vs. 0.13-0.21 for all other methods). This divergence is captured by the full R_NX_(K) profile, which reports invariance continuously across neighbourhood sizes. For three representative datasets spanning droplet-, spatial-, and plate-based technologies under a mild perturbation (downsampling to 95% of original counts), scFoundation and SCimilarity retained high RNX across nearly all neighbourhood sizes on the droplet- and plate-based datasets, declining only for the spatial dataset (Fig. 3a). In contrast, Geneformer’s score was markedly reduced at small neighbourhood sizes across all three datasets, indicating low local invariance, but increased steadily with neighbourhood size to approach the other methods at larger scales. This was particularly pronounced for the plate-based dataset, where RNX remained below 0.15 until neighbourhoods encompassed roughly half the dataset. Together, these profiles show that neighbourhood invariance distinguishes between preservation of global and local neighbourhoods, revealing representation properties that a single summary score would obscure.

**Figure 3.**
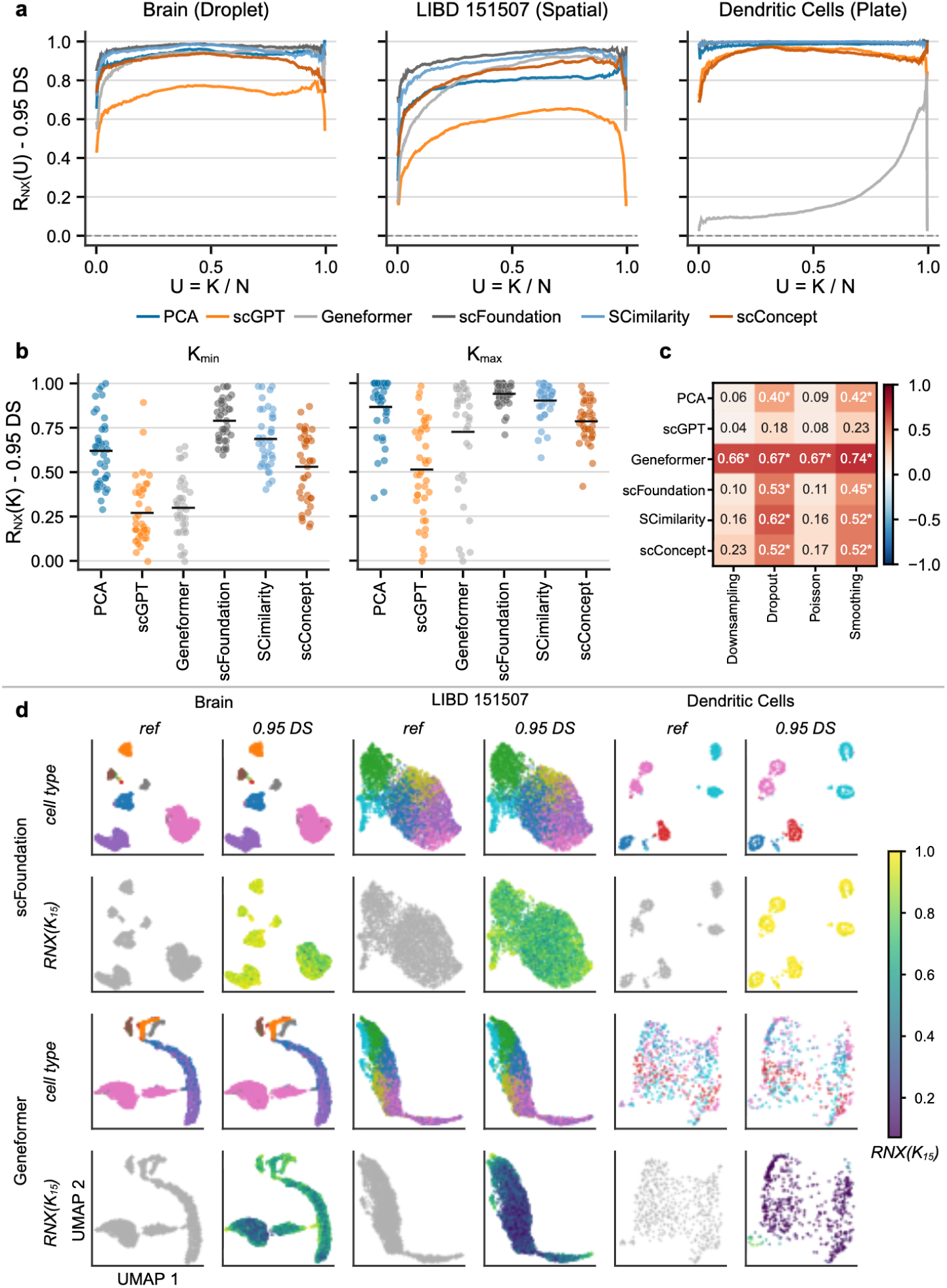
Geneformer preserves global structure while distorting local neighbourhoods. **(a)** R_NX_ scores as a function of neighbourhood size, expressed as the fraction U = K/N of the dataset, under a mild perturbation (downsampling to 95% of counts). Panels display three representative datasets spanning droplet-, spatial-, and plate-based technologies; each line is one embedding method. **(b)** R_NX_ evaluated at the smallest and largest neighbourhood sizes across all 39 datasets. Each point represents one dataset, and bars indicate the mean. **(c)** Spearman correlation between the number of annotated cell types in a dataset and perturbation-averaged Local R_NX_, computed across datasets separately for each embedding method and perturbation. Asterisks mark p < 0.05. **(d)** UMAP embeddings of the reference and the 95% downsampling perturbation, for scFoundation and Geneformer on the three datasets in (a). Rows alternate between colouring by annotated cell type and colouring by per-cell neighbourhood preservation at K=15. Reference panels remain grey in the preservation rows because preservation is defined exclusively for a perturbed embedding relative to its reference.

This scale dependence generalized across all 39 datasets (Fig. 3b). At the smallest neighbourhood size, Geneformer (0.30) and scGPT (0.27) showed the lowest neighbourhood structure invariance, trailing scFoundation (0.79), SCimilarity (0.69), PCA (0.62), and scConcept (0.53). At the largest neighbourhood size, however, Geneformer recovered to 0.73, approaching PCA (0.87) and scConcept (0.79), while scGPT remained the most distorted (0.51). The difference between R_NX_ at the smallest and largest neighbourhood sizes was 0.43 for Geneformer, compared to just 0.15-0.26 for all other methods. This pattern persisted across perturbations. Under Poisson resampling, for example, Geneformer ranked last at the smallest neighbourhood size but recovered to second at the largest (Fig. S10).

Geneformer’s scale dependence was also reflected in its relationship with dataset composition. Across all perturbations and metrics, its neighbourhood structure invariance scores correlated strongly with the number of annotated cell types in the dataset (Fig. 3c; Spearman ρ = 0.66-0.74, all p < 0.05). The effect was particularly pronounced under Poisson resampling, which most closely approximates pure technical noise. Under this perturbation, all other methods exhibited small, non-significant correlations (0.09-0.17), whereas Geneformer maintained a strong positive correlation (0.67). Because datasets containing more cell types generally exhibit greater large-scale transcriptional diversity, this observation is consistent with Geneformer preferentially preserving global rather than local neighbourhood structure.

A per-cell Uniform Manifold Approximation and Projection (UMAP) visualization illustrates how neighbourhood preservation varies across the embedding space (Fig. 3d). When coloured by cell type under a mild perturbation (downsampling to 95% of original counts), the reference and perturbed embeddings for both scFoundation and Geneformer appear similarly organized, with annotated cell types clustering together as expected. However, colouring cells by their local neighbourhood invariance score shows a different picture. scFoundation maintains high invariance uniformly across the embedding space, whereas Geneformer contains extensive regions of low neighbourhood preservation. The same visual organization that a standard analysis would take as evidence of a working embedding therefore coexists with substantial distortion of the local neighbourhoods that downstream analyses depend on.

One possible explanation for this scale dependence lies in Geneformer’s tokenization pipeline. It encodes each cell as a ranking of genes by normalized expression, discarding absolute expression magnitude. These rank transformations are inherently coarse; discarding magnitude eliminates fine-grained continuous variation, while preserving the larger differences that separate global clusters. However, we note that this interpretation is speculative and applies only to the checkpoint evaluated here (V1-10M); subsequent or larger checkpoints may behave differently.

### Neighbourhood preservation metrics track scIB bio-conservation scores

Current evaluations assess scFM embeddings against cell-type, batch, and trajectory labels^10–14,16,20^. We first summarize how the six methods perform under these reference evaluations (Fig. 4a,b). scIB bio-conservation scores separated the methods only weakly: five of the six fell within a narrow range (mean 0.55–0.58), with the refit PCA baseline (0.56) matching the scFMs and only Geneformer scoring lower (0.48). Batch correction reordered the methods substantially, with scConcept and scGPT ranking highest (mean rank 1.7 and 2.1) while PCA and scFoundation ranked lowest (5.4 and 4.8). This trade-off is well established in the design of scIB benchmarks^40^, and we confirmed that bio-conservation and batch-correction scores are moderately anticorrelated on the datasets and methods tested here (Fig. S11, r = −0.5, ρ = −0.4, p < 0.001). On trajectory inference (computed as the Spearman correlation between inferred and reference cell ordering), scFoundation ranked highest (mean rank 1.9) across the seven datasets tested. Model rankings therefore shift with the reference score chosen, and a simple PCA baseline remains competitive on bio-conservation, in line with the sensitivity of task-based comparisons noted previously.

**Figure 4.**
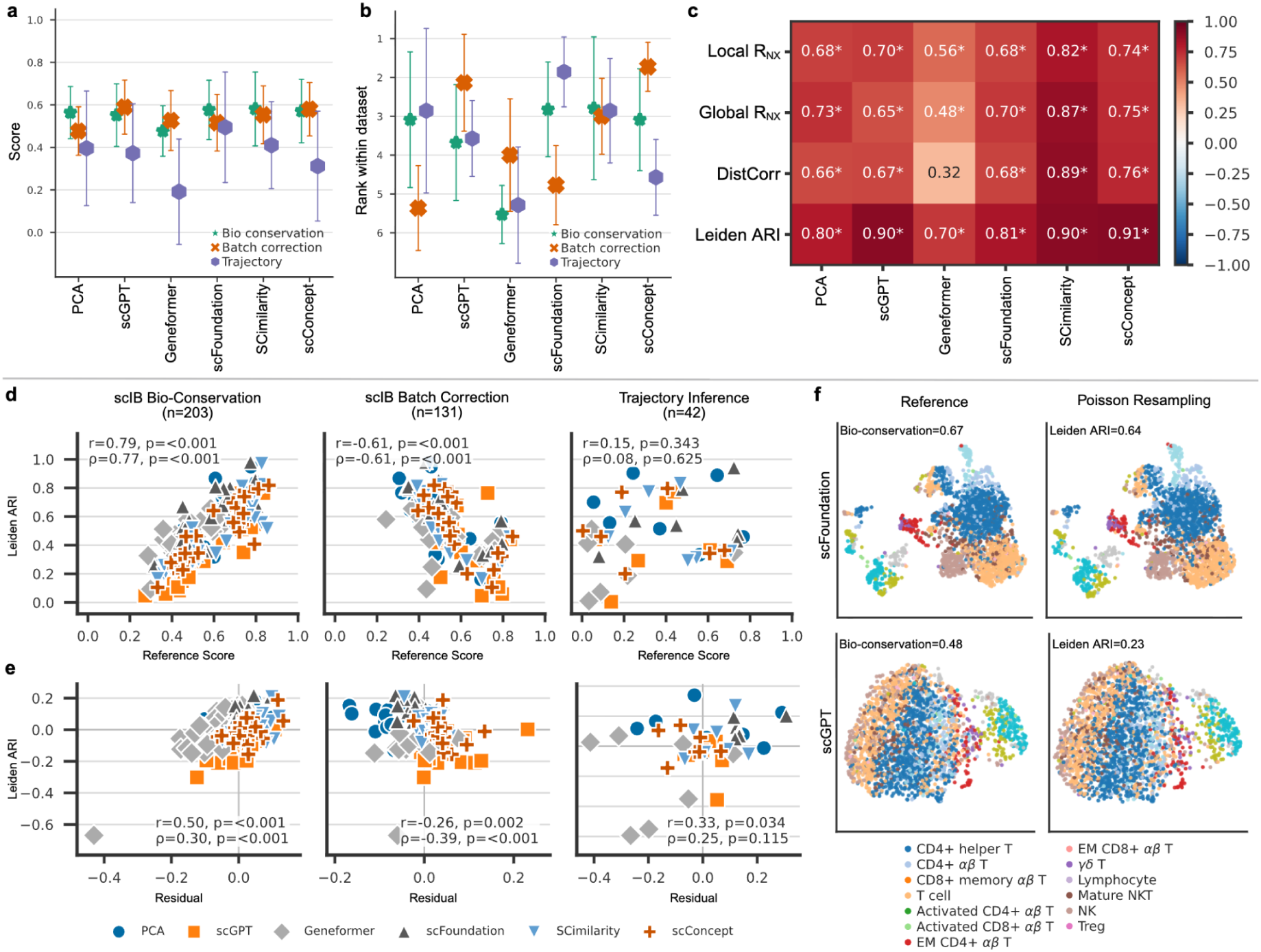
Label-free distortion metrics recapitulate scIB scores. **(a)** Mean ± s.d. reference scores across datasets for each embedding method, grouped by evaluation category: bio-conservation (green), batch correction (orange), and trajectory (purple). Bio-conservation is assessed on 34 cell-type tasks, batch correction on 22 batch tasks, and trajectory conservation on 7 trajectory tasks; each point is a per-dataset mean and error bars indicate the standard deviation across datasets within each category. **(b)** The same categories, with each point giving a method’s rank within a dataset (1 = best of the six methods). **(c)** Spearman correlation between each neighbourhood preservation metric (rows) and the scIB bio-conservation average, computed across the 34 cell-type datasets separately for each embedding method (columns). Values are annotated in cells, and asterisks mark p < 0.05. **(d)** Leiden ARI under a single Poisson-resampling iteration plotted against reference scores on three benchmarks: scIB cell-type conservation (left), scIB batch correction (centre), and trajectory (right). Each point is one dataset–method pair, with colour and marker denoting method. **(e)** As in (d), with each dataset-method pair shown as within-dataset residuals of both quantities (each centred on its dataset mean). **(f)** UMAP embeddings of a representative dataset (Breast T cells), both original and Poisson-resampling perturbation, for scFoundation and scGPT. Cells are coloured by annotated cell type.

We then asked how neighbourhood preservation metrics relate to these reference evaluations by correlating each metric with standard evaluation scores. Of the four neighbourhood preservation metrics, Leiden ARI tracked the reference scores most strongly, and we use it as the representative metric in Fig. 4d,e; the same pattern held for Local RNX, Global RNX, and DistCorr, more weakly and in the same directions (Fig. S12). Leiden ARI correlated positively with bio-conservation across all dataset-method pairs (r = 0.79, ρ = 0.78, n = 203) and negatively with batch correction (r = −0.61, ρ = −0.61, n = 131). This correlation with bio-conservation held within every method separately but varied in strength (Fig. 4c), it was strongest for SCimilarity (ρ = 0.82-0.89 across the four metrics) and weakest for Geneformer (ρ = 0.32-0.70, with its DistCorr correlation the only non-significant cell), while Leiden ARI was the most consistent metric across methods (ρ = 0.70–0.91 for every model). Trajectory inference showed no significant pooled association with Leiden ARI (r = 0.15, ρ = 0.08, p = 0.63, n = 42).

Pooled across datasets, the correlation could partly reflect differences in dataset difficulty rather than differences between methods. We therefore residualized both quantities against their dataset means, asking whether a method with greater neighbourhood preservation than other methods on a given dataset also scores higher on that dataset’s reference evaluation. The association persisted in the same directions, positive for bio-conservation (r = 0.50, ρ = 0.30) and negative for batch correction (r = −0.26, ρ = −0.39; Fig. 4e), although attenuated once dataset-level variation was removed. The correlation with trajectory inference remained weak, but was nominally significant by Pearson correlation (r = 0.33, p = 0.034) but not by rank correlation (ρ = 0.25, p = 0.12).

A concrete example illustrates this correspondence (Fig. 4f). On a breast T-cell dataset, whose closely related cell types make fine structure hard to preserve, scFoundation resolves more clearly separated structure in the reference embedding than scGPT, matching its higher bio-conservation score (0.67 versus 0.48). Under Poisson resampling, which adds sampling noise without changing cell identity, scFoundation also retains higher local neighbourhood invariance and a correspondingly higher Leiden ARI. The two scores move together because they depend on the same property: an embedding whose local neighbourhoods are biologically organized is also one whose neighbourhoods, and therefore whose Leiden partition, survive resampling. Where a representation lacks that structure, both scores fall together.

## Discussion

Six representations that current benchmarks rank closely differ substantially in how much of their structure survives a resampling of the data. Under a perturbation that retains 95% of counts, a level of variation no analyst would treat as material, Local RNX across the six ranged from 0.89 to 0.49, and the spread widened at finer scales rather than closing. Geneformer illustrates this most clearly: it preserved coarse organization comparably to the other representations (Global RNX 0.73) while retaining substantially less local structure (Local RNX 0.30), a gap more than twice that of any other method. This occurred despite visually coherent cell-type organization in the embedding, illustrating how conventional inspection can miss local instability (Fig. 3d). Analyses that depend on fine neighbourhood structure, such as resolving graded transitions or identifying rare populations, may therefore rest on structure that is sensitive to the particular sampling realization.

Standard evaluation does not directly measure this property. A benchmark score establishes that an embedding supports a particular task, but not whether the underlying representation is stable to nuisance variation in the measurements themselves. Our label-free scores correlate with scIB bio-conservation, consistent with the idea that both evaluations favour neighbourhood structure that reflects reproducible biological variation. Neighbourhood structure invariance therefore provides a complementary criterion for evaluating representations, asking whether the structure on which downstream analyses operate is stable before those analyses are applied. This distinction has practical consequences beyond model comparison. External annotations are often unavailable, incomplete, or difficult to define precisely in the settings where general-purpose representations are most useful. Because our evaluation requires only the expression data and a correspondence between perturbed and reference cells, it can be applied before downstream analysis and without task-specific labels. The same formulation also extends naturally to multimodal representations and other settings in which cells can be embedded but no single annotation adequately describes the joint space.

Neighbourhood structure invariance is not sufficient to define a good representation, since a representation that discarded biological information entirely could remain perfectly invariant, and we therefore view it as a complement to downstream measures of selectivity and biological utility rather than a replacement. The same perturbation-based framework could be extended in the opposite direction, using perturbations designed around biological programs or functional gene groups to test whether representations respond selectively to biologically meaningful changes, so that invariance to nuisance variation and selectivity to biological variation together provide a more complete characterization of representation quality. The narrower conclusion supported here is that representations appearing similarly successful under current benchmarks can differ markedly in the stability of the neighbourhood structure they provide, and we therefore suggest that neighbourhood structure invariance to nuisance perturbations should be measured alongside downstream task performance when developing and evaluating single-cell foundation models.

## Methods

### Evaluation framework

We evaluated single-cell foundation model representations by applying nuisance perturbations to raw count matrices, embedding both the unperturbed and perturbed matrices, and measuring the distortion of neighbourhood structure relative to the unperturbed reference. For each dataset, we prepared a reference count matrix *X* (cells x genes) and, for each perturbation *τ*, a perturbed matrix *X’* of identical shape. Both were embedded independently by the same model to generate *E* and *E’*, and all metrics compared the aligned embedding pairs *(E_i_, E_i_’)*, which correspond one-to-one by construction. All structure metrics were computed in embedding space.

### Nuisance perturbations

Each perturbation acts on the count matrix while preserving cell identity to maintain a one-to-one mapping between perturbed and unperturbed matrices.

- **Poisson resampling** redraws each non-zero count from Poisson(λ = count) using rng.poisson(), then repeats this resampling for t chained iterations (iteration i uses the counts from iteration i−1), injecting count noise while keeping the sparsity pattern of non-zeros. Tested over t ∈ {1, 2, 3, 4}.
- **Downsampling** applies binomial subsampling of counts at rate f by drawing rng.binomial(n = count, p = f) for each non-zero entry, emulating reduced sequencing depth. Tested over f ∈ {0.4, 0.6, 0.8, 0.9, 0.95}.
- **Gene dropout** sets each non-zero entry to zero with probability d by drawing rng.random() independently for each non-zero entry and zeroing those below the dropout threshold, emulating incomplete transcript capture. Tested over d ∈ {0.1, 0.2, 0.4, 0.6}.
- **Local smoothing** replaces each cell’s profile with a k-nearest-neighbour average. We log-normalized a copy of the input for neighbour finding only (scanpy.pp.normalize_total(…, target_sum = 10,000) followed by scanpy.pp.log1p() and scanpy.pp.pca(…, n_comps = 50, svd_solver = "arpack")), built a kNN graph with sklearn.neighbors.NearestNeighbors(n_neighbors = k) in PCA space, and averaged raw counts over k neighbours including self with uniform weights 1/k. Smoothed counts were rounded to the nearest integer with numpy.rint(). Tested over k ∈ {5, 10, 20, 50, 100}.

### Distortion metrics

For each perturbation, we computed four metrics between the reference and perturbed embeddings, each ranging from -1 to 1, with higher values indicating greater preservation.

- **DistCorr**, as defined by Székely,^35^ measures correlation between the pairwise-distance matrices of *E* and *E’*, with double-centring. We computed Euclidean pairwise distances with scipy.spatial.distance.pdist, converted them to square matrices with scipy.spatial.distance.squareform, double-centred each matrix (row, column, and global means removed), and calculated DistCorr from the inner product of the centred matrices normalized by their respective variances.
- **Local R_NX_** and **Global R_NX_** are the local and global structure-preservation scores as implemented in ViScore^38^, derived from the co-ranking matrix of *E* and *E’*, following the R_NX_ formulation of Lee and Verleysen^36^. We called viscore.score(E_ref, E_p) and recorded score["Sl"] as Local RNX, score["Sg"] as Global RNX, and the full score["RNX"] curve over neighbourhood sizes k = 1, …, n−1.
- **Leiden ARI** is the adjusted Rand index^39^ between independent Leiden clusterings of *E* and *E’*. For each resolution ∈ {0.25, 0.5, 1.0, 2.0}, we built a k = 15 nearest-neighbour graph with Euclidean distance using scanpy.pp.neighbors(…, n_neighbors = 15, metric = "euclidean"), ran Leiden clustering separately on each embedding with scanpy.tl.leiden(…, flavor = "igraph", random_state = seed), and computed sklearn.metrics.adjusted_rand_score(labels_ref, labels_p). Reported values are the mean ARI across resolutions.

### Permutation nulls

For each dataset, model, and perturbation setting, we computed a null distribution for the DistCorr metric by randomly shuffling the correspondence between reference and perturbed cells and recomputing the metric over 100 permutations. We pre-computed the double-centred distance matrices; for each permutation b = 1, …, 100, drew a cell permutation π with numpy.random.Generator.permutation(n), permuted the manipulated distance matrix accordingly, and computed null DistCorr_b. Reported values are corrected against the null mean μ_null = mean(null DistCorr_b) as null_corr = (obs − μ_null) / (1 − μ_null)

### Reference benchmark metrics

Label-based reference metrics were computed on the reference embeddings only, without downsampling. When applicable, cell type, batch, and trajectory metrics were computed using the dataset’s original labels. For a full list of datasets and available labels, see supplementary data S1.

scIB metrics were computed using the scib-metrics Python package^40,43^. For each reference embedding E_ref, we ran scib_metrics.benchmark.Benchmarker with BioConservation(isolated_labels, silhouette_label, clisi_knn, nmi_ari_cluster_labels_leiden) and BatchCorrection(bras, ilisi_knn, kbet_per_label, graph_connectivity). Bio-conservation score was calculated as the mean of isolated_labels, silhouette_label, clisi_knn, and the Leiden NMI and ARI; batch correction as the mean of bras, ilisi_knn, kbet_per_label, and graph_connectivity.

Trajectory inference was assessed on reference embeddings for the trajectory-benchmark datasets by diffusion pseudotime. We computed a kNN graph (k = 15) and diffusion map (10 components) with scanpy.pp.neighbors and scanpy.tl.diffmap, set the DPT root to the cell nearest the centroid of the earliest milestone, and obtained pseudotime with scanpy.tl.dpt. The reported score is the absolute Spearman correlation between pseudotime and curated milestone order from scipy.stats.spearmanr. For null calibration, we shuffled milestone labels with numpy.random.Generator.permutation(reference_order) over 10 permutations and recomputed the Spearman score to obtain an empirical null mean and p-value.

### Sample-size analysis

To determine whether distortion metrics are stable under subsampling, we ran an experiment on six human atlases (arterial, brain, immune, lung, retina, Tabula Sapiens), embedding with SCimilarity^41^. For each atlas and target subsample size n ∈ {200, 500, 1000, 2000, 5000, 10000, 20000}, we drew five independent uniform random subsamples using seeds 0-4. Subsampling was performed with pandas.DataFrame.sample(n = n, random_state = seed). Each subsample was then run through the same pipeline as the main benchmark: the full nuisance-perturbation grid from configs/default.yaml.

For each atlas, subsample size, perturbation type, perturbation parameter, and metric, we formed a stability curve over n. At each n, we computed the seed-averaged metric (mean across the five subsample seeds) and the cross-seed coefficient of variation CV = std / |mean| using pandas groupby aggregation with sample standard deviation. To summarize precision at each atlas and subsample size, we took the median CV across all perturbation curves and estimated 95% confidence intervals on that median by bootstrap (10,000 resamples with numpy.random.default_rng(0)). We used CV ≤ 0.05 as a precision threshold in the stabilization plots. Convergence was defined per curve as the smallest subsample size n beyond which every subsequent increase in n changed the seed-averaged metric by at most 0.05 in absolute value; curves that did not meet this criterion within the evaluated range were assigned a convergence size of infinity and reported as the maximum evaluated subsample size (20,000 cells).

### Embeddings using foundation models and PCA baseline

Embeddings were generated using our custom repository, transcriptomic-fms, located at https://github.com/ahmedh-sallam/transcriptomic-fms. Compute used Apptainer containers on H100 GPU. Each model was run in a dedicated container with fixed weights.

- **PCA:** log-normalization (10,000 counts per cell, log1p), scaling, then PCA (sklearn.decomposition.PCA, svd_solver = "arpack", seed 42) with 128 components.
- **scGPT:** Pretrained human scGPT; HVGs subset using scanpy.pp.highly_variable_genes(flavor="seurat_v3"), expression binned (51 bins) via the scGPT preprocessor, cell embeddings generated from the last hidden layer. scGPT used the pretrained human scGPT checkpoint (scGPT_human). Non-zero counts were binned into 51 expression bins (default n_bins = 51) without additional normalization in the scGPT preprocessor, and cell embeddings were generated with scgpt.tasks.embed_data(…, gene_col = "gene_symbols", device = cuda) from the model’s cell-representation layer (X_scGPT). Cells with no vocabulary overlap or all-zero expression after gene filtering were dropped before embedding.
- **Geneformer:** Geneformer V1-10M (ctheodoris/Geneformer, subdirectory Geneformer-V1-10M). Cells were tokenized with geneformer.TranscriptomeTokenizer using the GC30M token dictionary, rank-value encoding, model_input_size = 2048, and special_token = False. Tokenized cells were passed to geneformer.EmbExtractor in cell mode with emb_layer = -1 (final transformer layer). Embedding rows were realigned to the input cells using the preserved cell_id field after Geneformer’s internal length sorting.
- **scFoundation:** 01B-resolution checkpoint in cell-embedding mode (version = ce, output_type = cell, pool_type = all). Gene symbols were uppercased and mapped to scFoundation’s fixed 19,264-gene human index (OS_scRNA_gene_index.19264.tsv), padding missing genes with zeros. Raw counts were converted to log1p(CPM) within each cell (pre_normalized = F), augmented with target high-resolution tokens tgthighres = t4 and log10 total-count tokens.
- **SCimilarity:** v1.1, Raw counts were aligned to the model gene order with scimilarity.utils.align_dataset after scimilarity.utils.consolidate_duplicate_symbols, log-normalized with scimularality.utils.lognorm_counts, and embedded with scimilarity.CellEmbedding.get_embeddings.
- **scConcept:** Corpus-30M contrastive model; Ensembl IDs and raw counts passed to extract_embeddings. Cell embeddings were extracted with concept.scConcept.extract_embeddings(…, gene_id_column = "gene_id"), retaining the cls_cell_emb output as the cell representation.

## Data availability

We used 39 datasets spanning cell-type integration, batch correction, trajectory inference, and spatial transcriptomics (Data S1). These were pre-curated by multiple evaluations^10,44–46^. All datasets are publicly available and were retrieved from their primary repositories (See Data S1). All datasets were normalized to a common format: filtering to protein-coding genes, and counts taken from raw.X or count layers. Datasets flagged for subsampling were subsampled to 5,000 cells after preprocessing.

Ten datasets were downloaded from CZ CELLxGENE Discover^47^, listed along with their dataset version UUID: brain (a1ccad25-1438-41f6-a09e-d4960ca36b67) and spinal cord (72dab565-a96b-4b06-a1b7-3413efeff523)^48^, liver CRLM–NMP atlas (a3bcb68b-a36d-460f-bdde-da4fc0771c8c)^49^, breast B cells (90377993-524f-40d2-8295-6317288ffca6) and T cells (bdc0daca-3fcf-4c05-9d46-330e038b820c)^50^, large intestine (681f6f1b-f9b6-4d1d-8588-97fedad51f7a)^51^, developing entorhinal cortex (690cd774-112b-489d-af6c-8fd8aa2de80b)^52^, ileum(a5d350b0-a7a9-411c-b120-6ad57a26cd22)^53^, adult kidney (dad69d7e-fb52-46cd-88bc-e4926efcacce), and Tabula Sapiens stomach (3e8bd2b2-fe53-4b95-abb2-ce971b4f1c14)^54^. Six datasets were obtained from GEO: the A549 TGF-β1 epithelial-to-mesenchymal transition time course (GSE147405)^55^, stem-cell-derived β-cell differentiation (GSE114412)^56^, murine haematopoietic stem and progenitor cells following in vivo IFN-α treatment (GSE226824)^57^, the CellBench cell-line mixture benchmark (GSE118767)^58^, blood dendritic cells and monocytes (GSE94820)^59^, and IFN-β-stimulated peripheral blood mononuclear cells (GSE96583)^60^. Seven datasets were obtained from Zenodo: Human germline, mesoderm development and the cell cycle, were taken from the dynverse collection of single-cell omics datasets containing a trajectory (10.5281/zenodo.1443566)^46^. HER2-positive breast spatial transcriptomics sections for patients E–H were taken from the HER2ST deposition (10.5281/zenodo.4751624)^61^, with the corresponding cluster labels obtained from the accompanying repository at https://github.com/almaan/her2st (10.5281/zenodo.5511762). Two datasets were obtained from figshare (10.6084/m9.figshare.12420968): human immune atlas and the pancreas complexBatch task^40^. The embryoid body scRNA-seq time course was obtained from Mendeley Data (10.17632/v6n743h5ng.1)^7^. The 68k PBMC reference^62^ was obtained from the 10x Genomics single-cell gene expression datasets portal. Twelve Visium sections of human dorsolateral prefrontal cortex^63^ were retrieved through the Bioconductor package spatialLIBD^64^ using spatialLIBD::fetch_data(type = "sce"): 151507, 151508, 151509, 151510, 151669, 151670, 151671, 151672, 151673, 151674, 151675 and 151676.

## Code availability

Software analyses used Python 3.11.14 with pandas v2.3.3, scanpy 1.11.5, anndata 0.12.13, scib-metrics 0.5.9, ViScore 0.0.2 (commit eca4be7), scikit-learn 1.8.0, and umap-learn 0.5.12, as pinned in the project uv.lock. All code for perturbation and metrics implementation, analysis, and visualization dashboard is available through GitHub at https://github.com/ahmedh-sallam/scfm-controlled-manipulations.

For easier visualization of raw metrics, we deployed all raw metrics across all perturbations using a streamlit dashboard, available at: https://scfm-controlled-manipulations.streamlit.app/.

## Acknowledgments

We thank Dr. Leon French and Dr. Peter Koo for helpful discussions and conversations.

