## Supplementary Material for "Robustness to nuisance perturbations enables unsupervised evaluation of single-cell foundation models"

#### Supplementary Figures

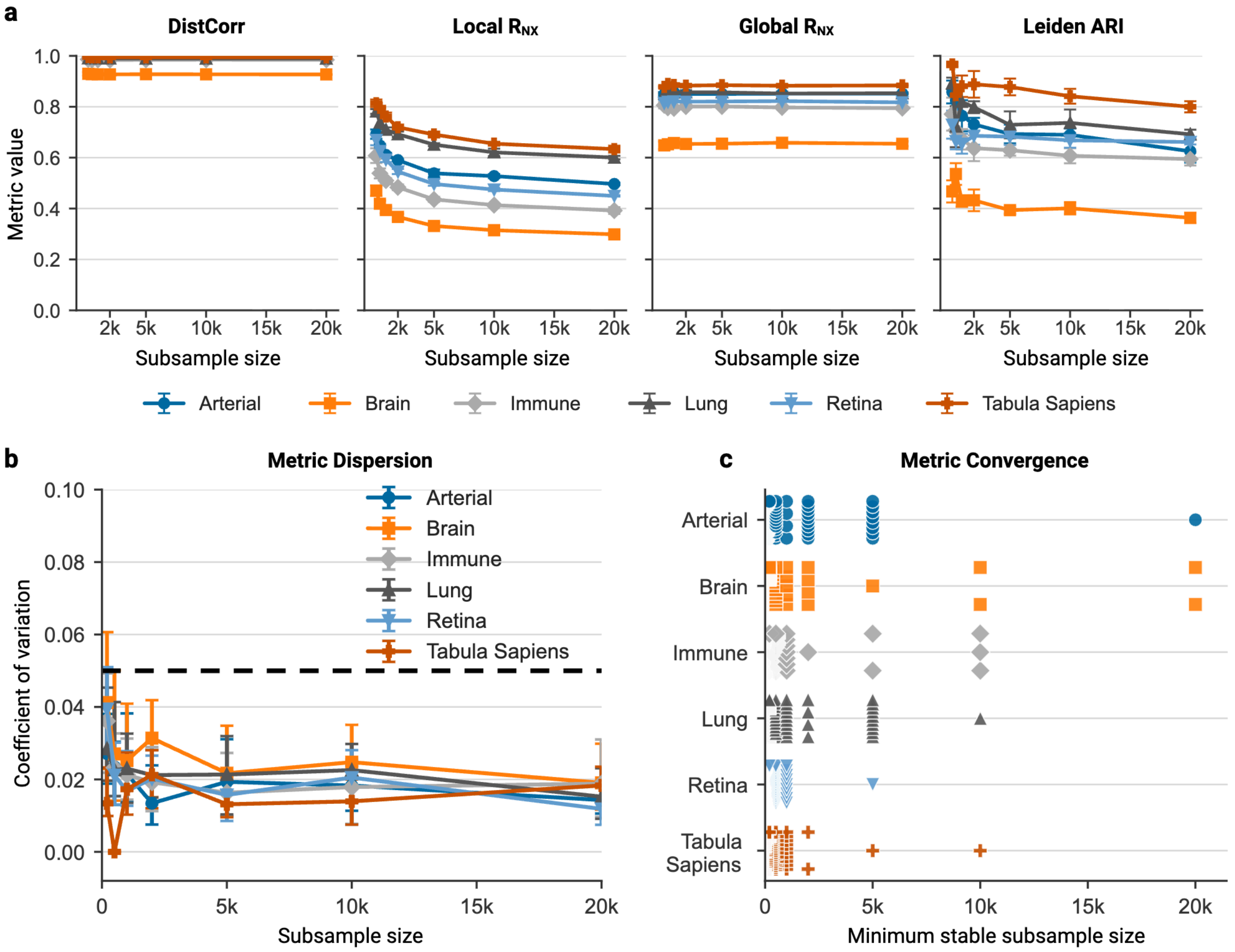


**Figure S1.** Subsampling validation. For each atlas, random subsamples were drawn across a range of sizes with five random seeds per size. Each subsample was subjected to 23 perturbations and embedded using SCimilarity. **(a)** Sample metric values against subsample sizes for one perturbation, one panel per metric, coloured by atlas. Points show the seed-averaged value; error bars, 1 std across seeds. **(c)** Dispersion. For each atlas, the median coefficient of variation (CV) across seeds, summarized over all perturbations and metrics at each subsample size. CV is the standard deviation of seed-level metric values divided by their absolute mean. Error bars, 95% CI across the summarized curves. Dashed line, the CV = 0.05 precision threshold. **(d)** Convergence. Distribution of stabilization subsample sizes within each atlas, one point per perturbation and metric combination. The stabilization subsample size is the smallest size beyond which every further increase changes the seed-averaged metric by at most 0.05 in absolute value.

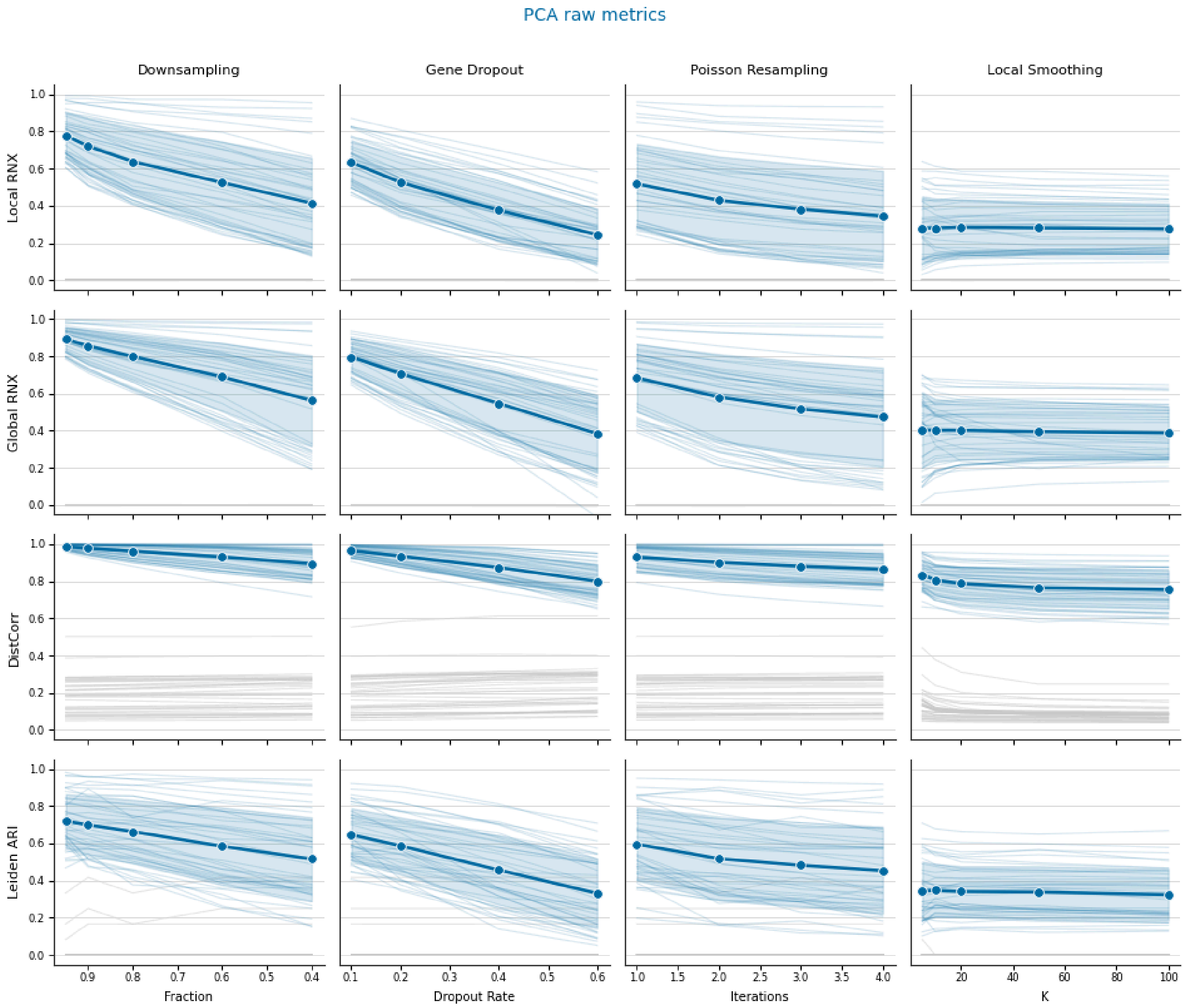


**Figure S2. Per-dataset perturbation grid, PCA.** Structure preservation as a function of perturbation strength, with rows corresponding to metrics and columns to perturbations. Each faint line is one of the 39 datasets, and gray lines are mean permutation nulls per dataset. Dark line shows cohort mean and shaded area is 1 std.

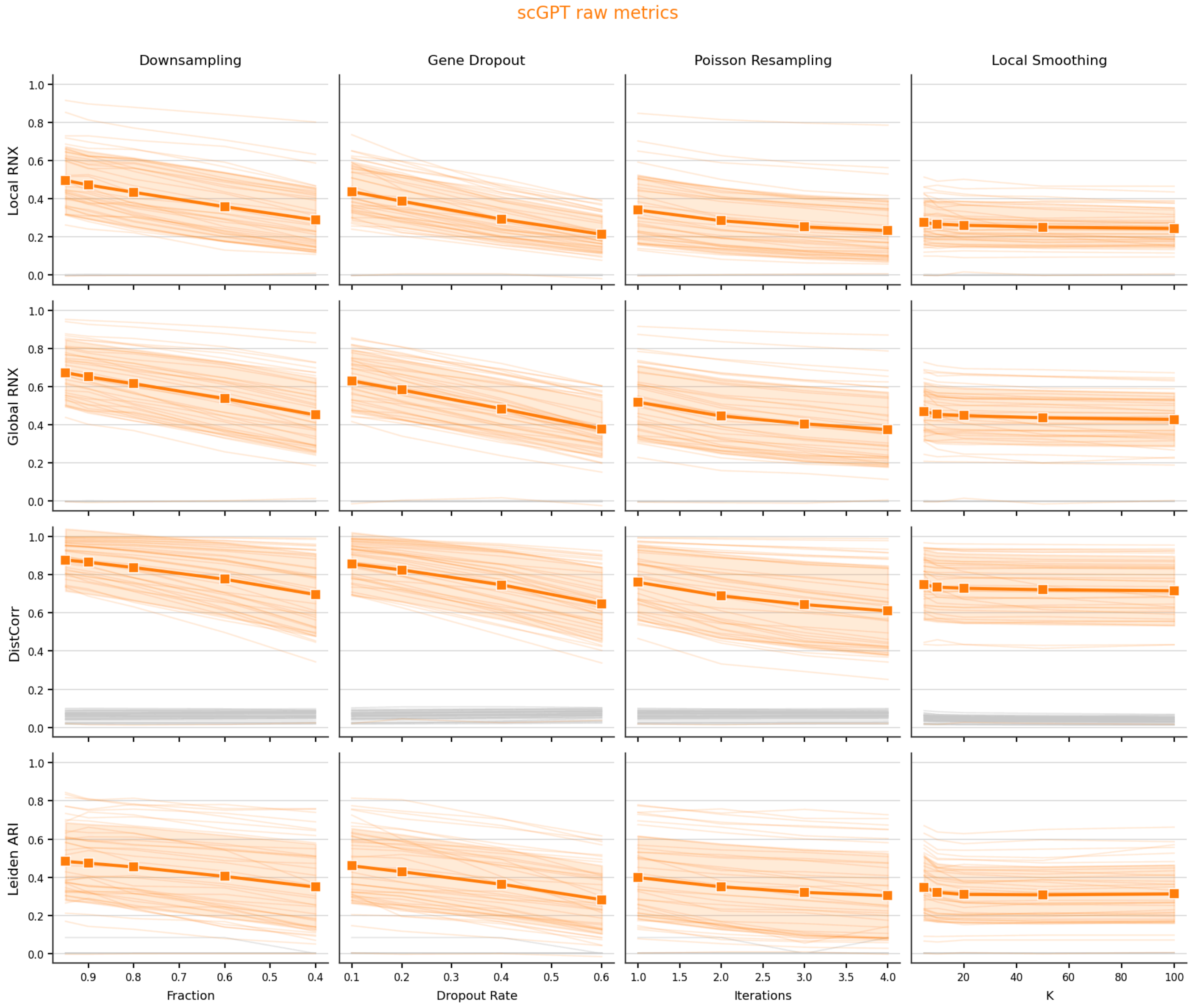


**Figure S3. Per-dataset perturbation grid, scGPT.** Conventions as in Figure S2.

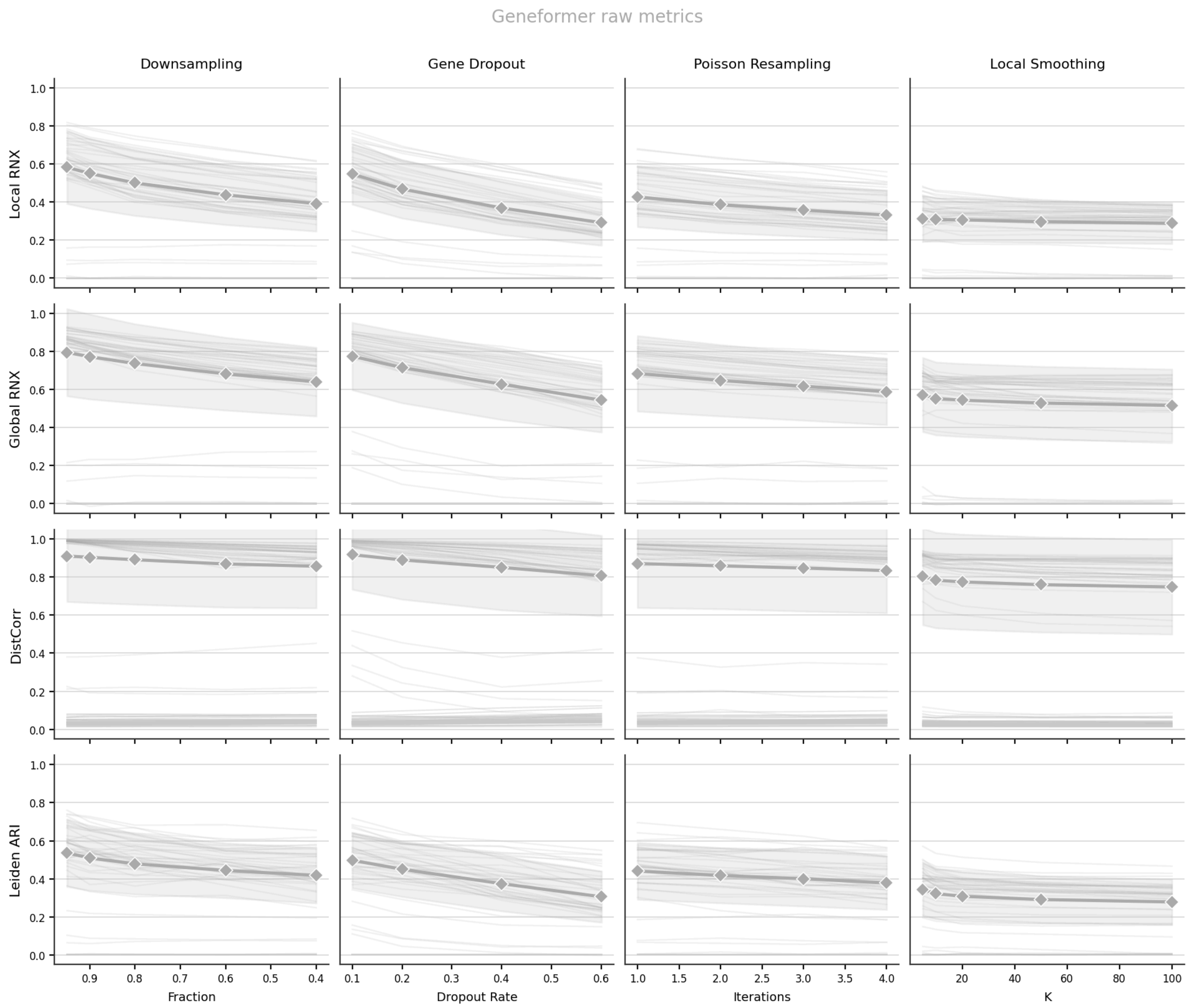


**Figure S4. Per-dataset perturbation grid, Geneformer.** Conventions as in Figure S2.

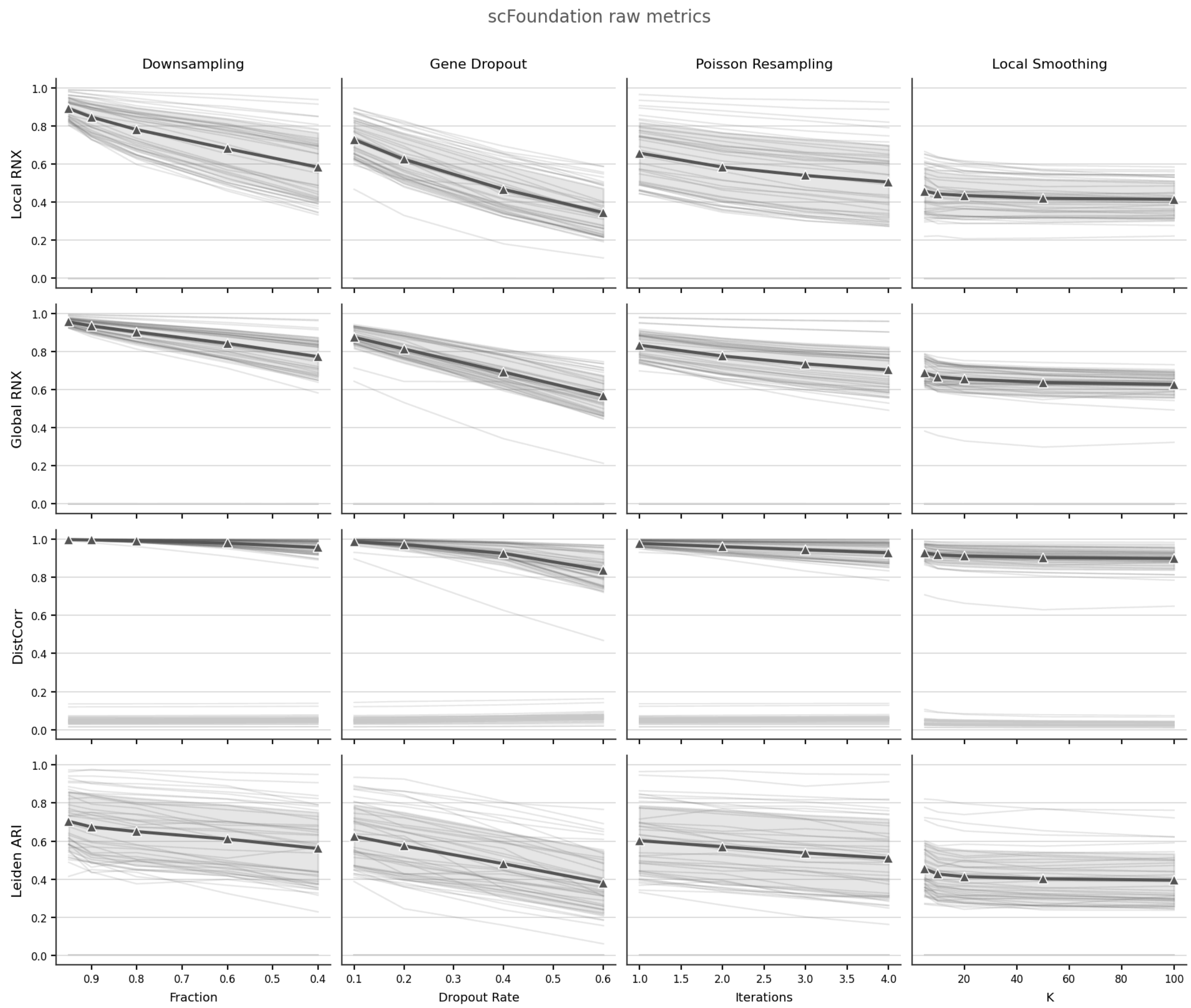


**Figure S5. Per-dataset perturbation grid, scFoundation.** Conventions as in Figure S2.

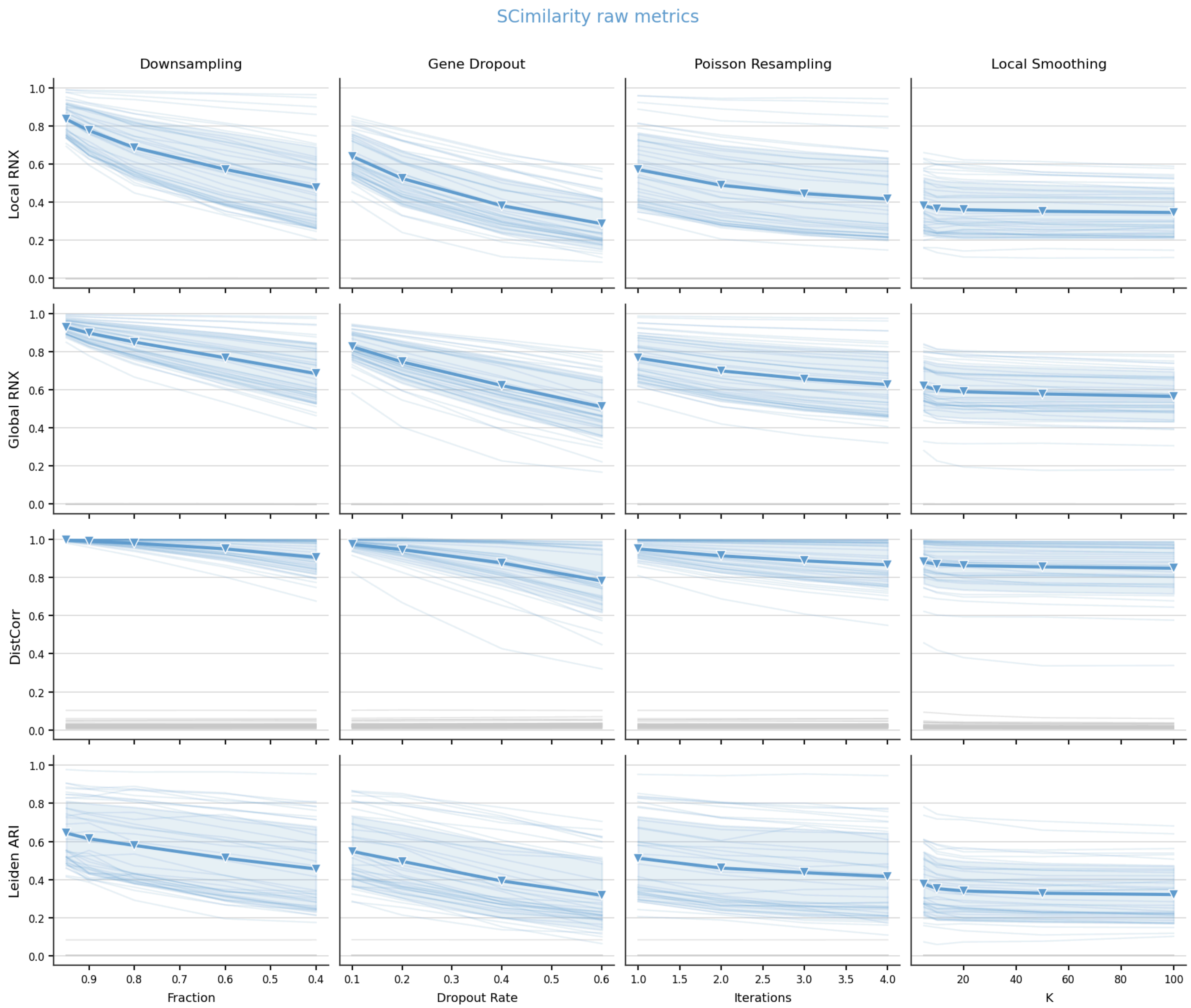


**Figure S6. Per-dataset perturbation grid, SCimilarity.** Conventions as in Figure S2.

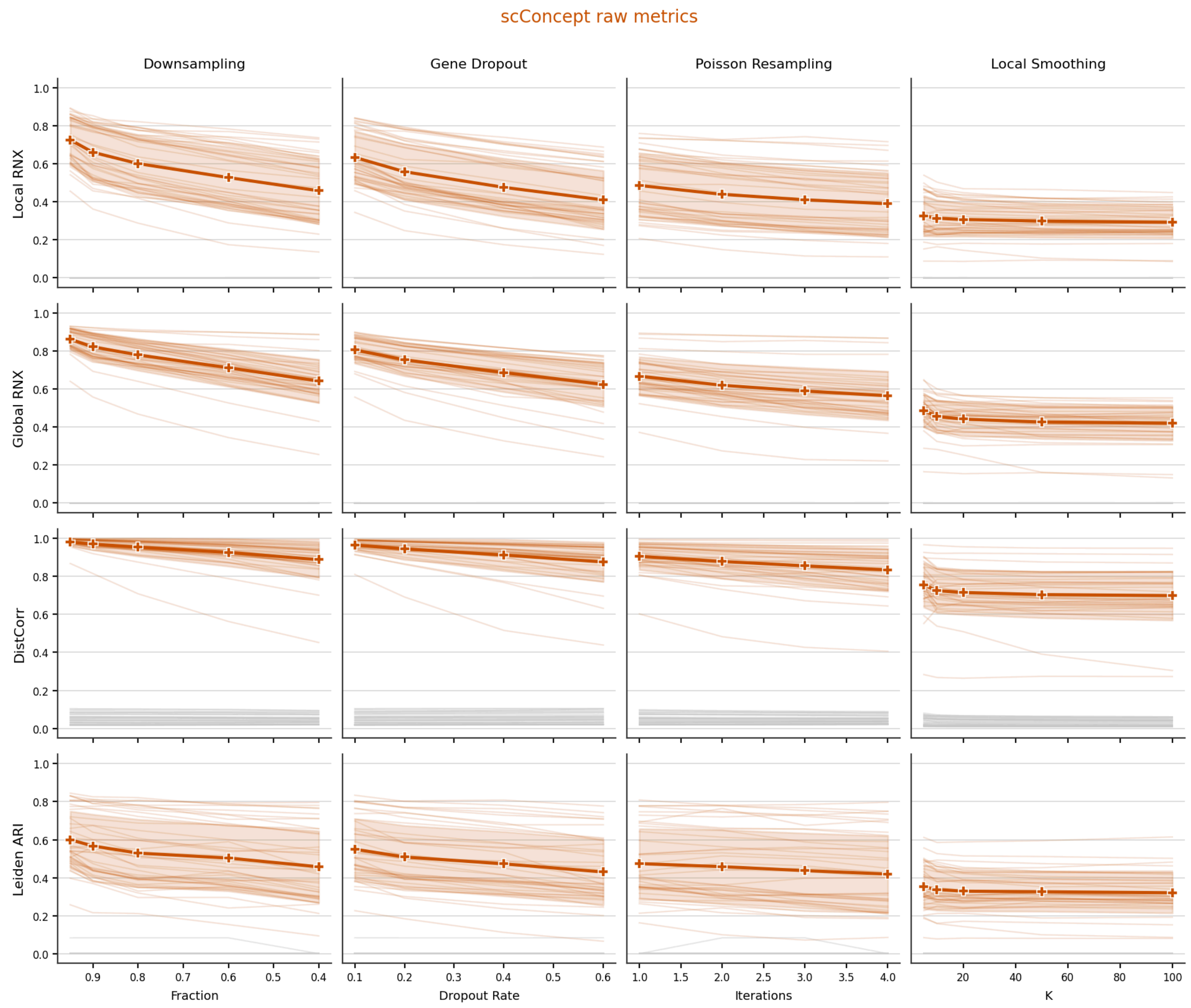


**Figure S7. Per-dataset perturbation grid, scConcept.** Conventions as in Figure S2.

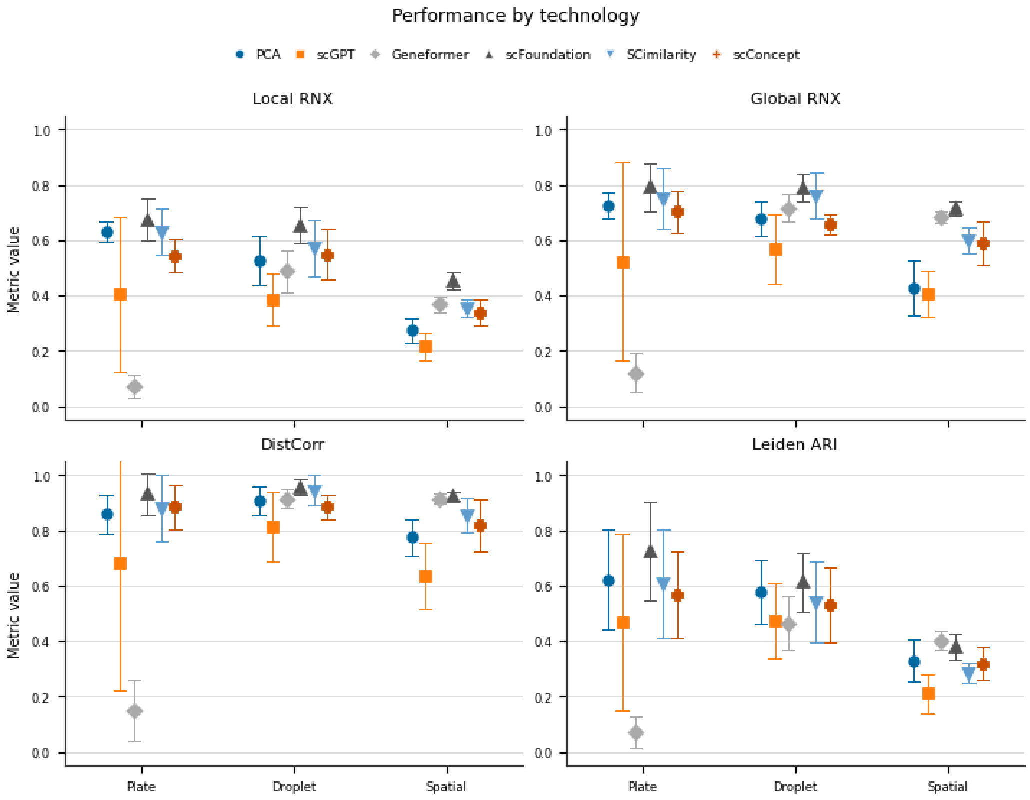


**Figure S8. Score by assay technology**, averaged over perturbations. Points represent the mean per representation and technology group, with error bars denoting ±1 SD.

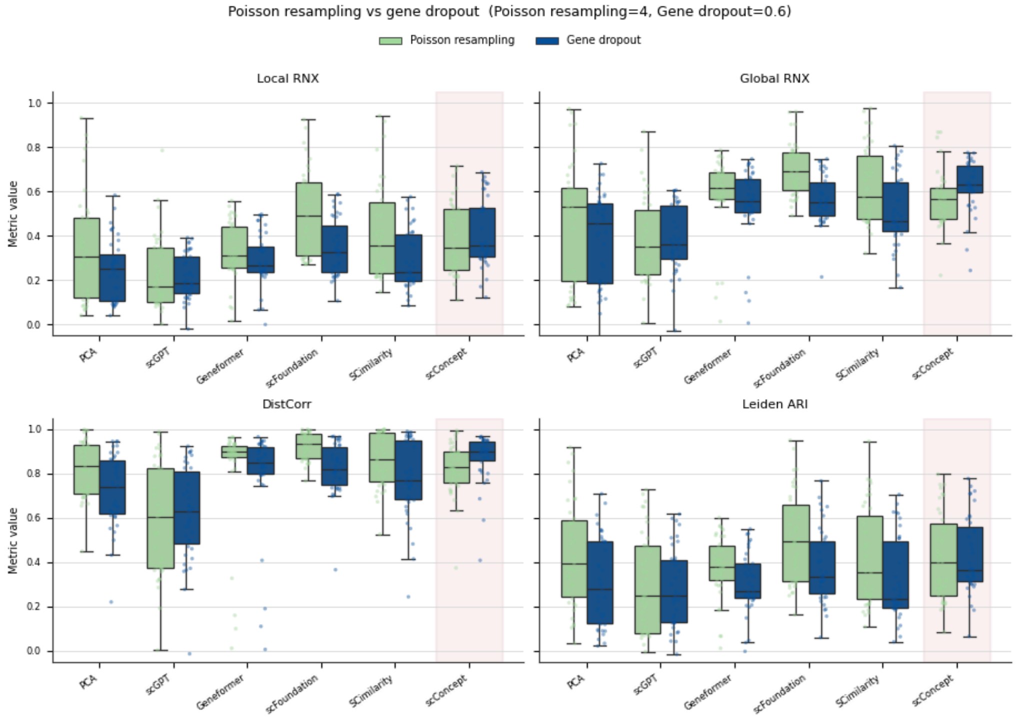


**Figure S9.** All metrics under Poisson resampling (iterations=4) and gene dropout (fraction=0.6), with one point per dataset. scConcept is highlighted as the model preserving the most global structure under gene dropout. The center line corresponds to the median, with boxes indicating the interquartile range (IQR) and whiskers extending to 1.5× IQR.

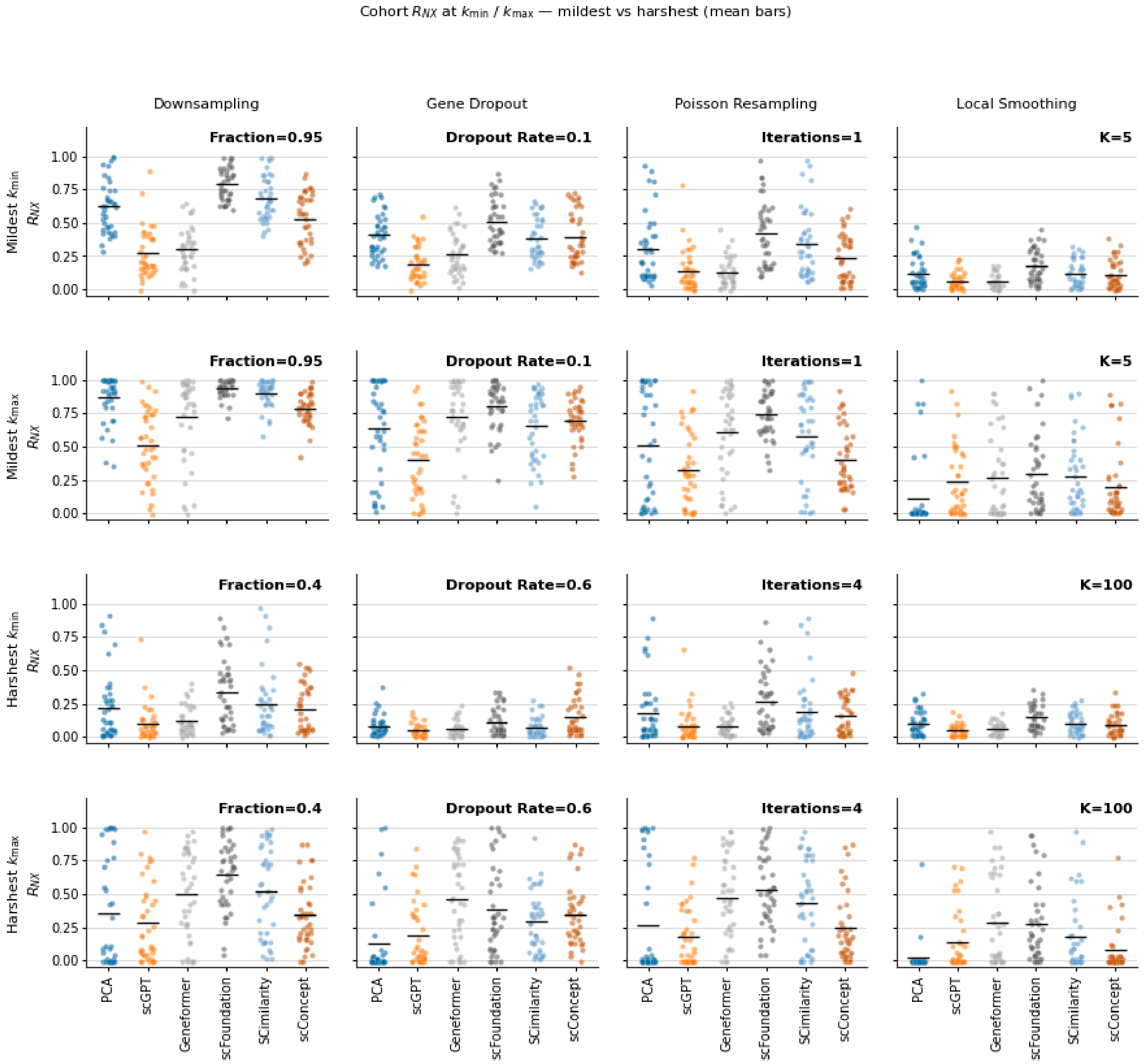


**Figure S10. Cohort RNX at neighbourhood extremes under the mildest and harshest perturbations.** Each panel shows raw embedding-space $R_{NX}$ across datasets with the cohort mean (black bar). Columns are perturbations (downsampling, gene dropout, Poisson resampling, local smoothing); rows are mildest vs harshest parameter settings at $k_{min}$ (local) and $k_{min}$ (global). The perturbation strength used in each panel is annotated at the top right (for downsampling, a higher fraction = milder; for the others, a larger parameter = harsher).

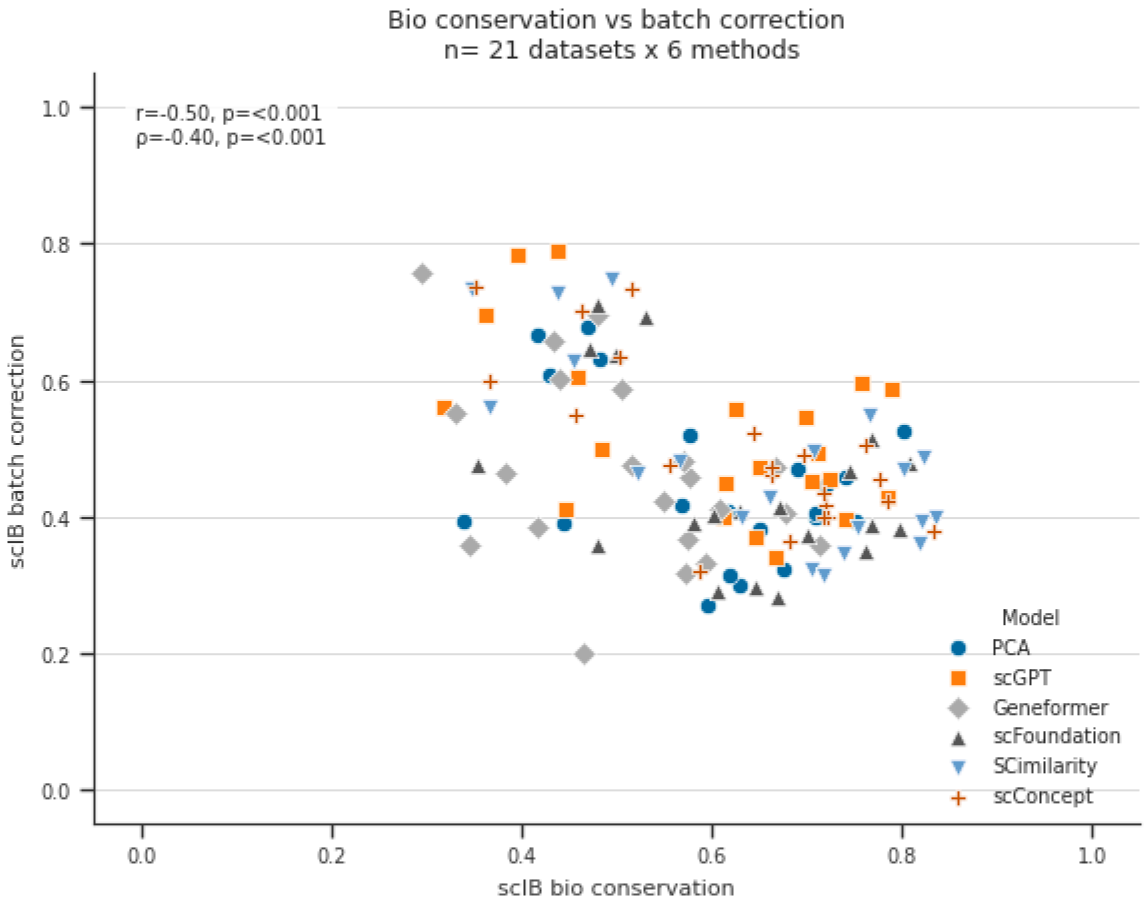


**Figure S11. Relationship between scIB bio-conservation and batch correction across embedding models.** Each point is one model×dataset pair (125 points; 21 curated datasets that provide both cell-type and batch labels). The x-axis is the scIB bio-conservation score and the y-axis is the scIB batch-correction score on the reference embedding. Points are coloured and shaped by model (PCA, scGPT, Geneformer, scFoundation, SCimilarity, scConcept). Pearson r and Spearman ρ (inset) summarise the association across all model×dataset points.

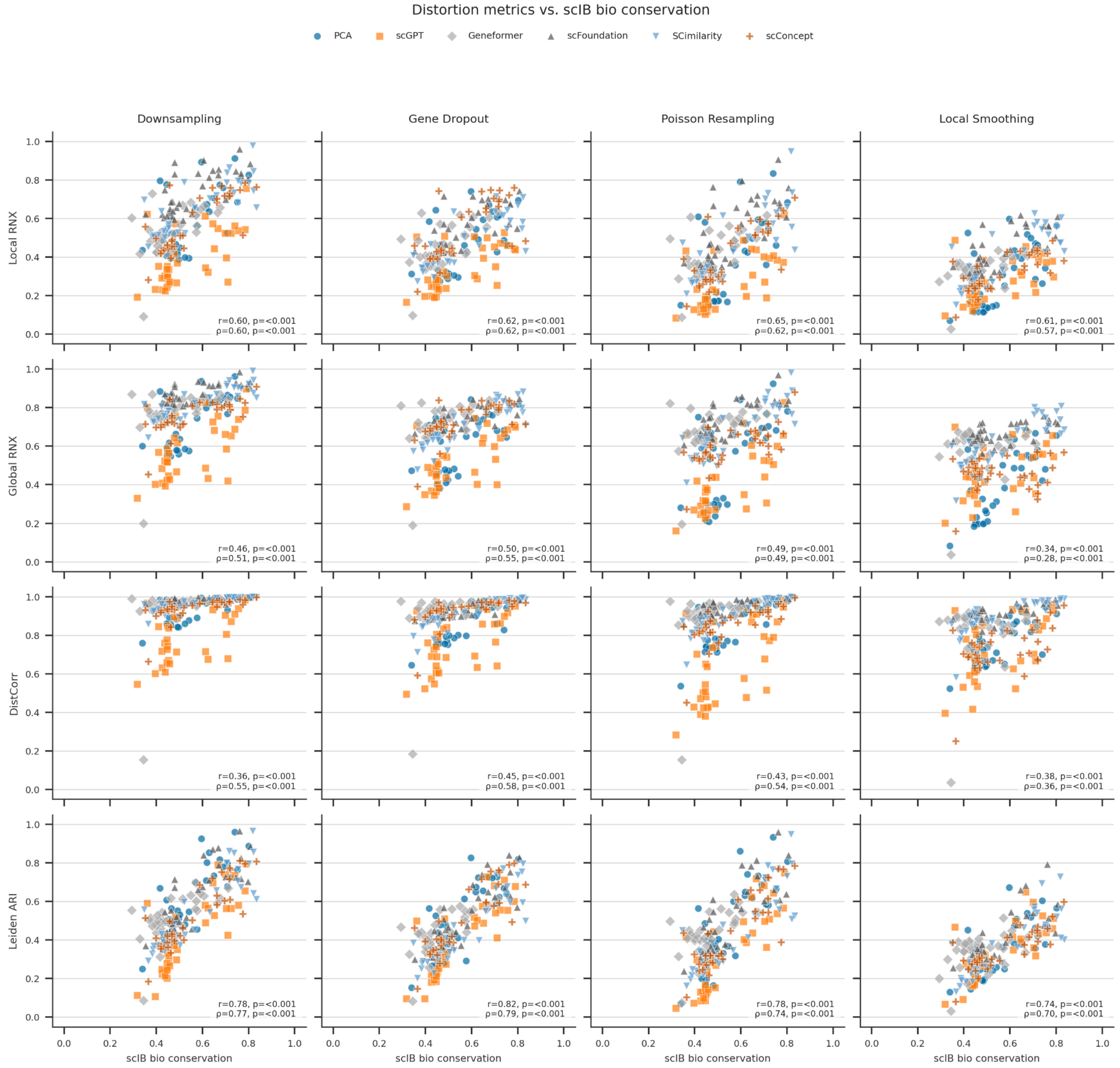


**Figure S12. Label-free distortion metrics vs. scib bio conservation score.** Each panel shows one distortion metric (rows: Local RNX, Global RNX, DistCorr, Leiden ARI) under one controlled perturbation (columns: downsampling, gene dropout, Poisson resampling, local smoothing). The y-axis is the null-corrected distortion score, averaged over perturbation strengths; the x-axis is the mean of scIB bio-conservation metrics on the reference embedding. Each point is one model×dataset pair (datasets with cell-type labels); colour and marker encode the embedding model (PCA, scGPT, Geneformer, scFoundation, SCimilarity, scConcept). Insets report Pearson r and Spearman ρ with p-values across all model×dataset points in the panel.

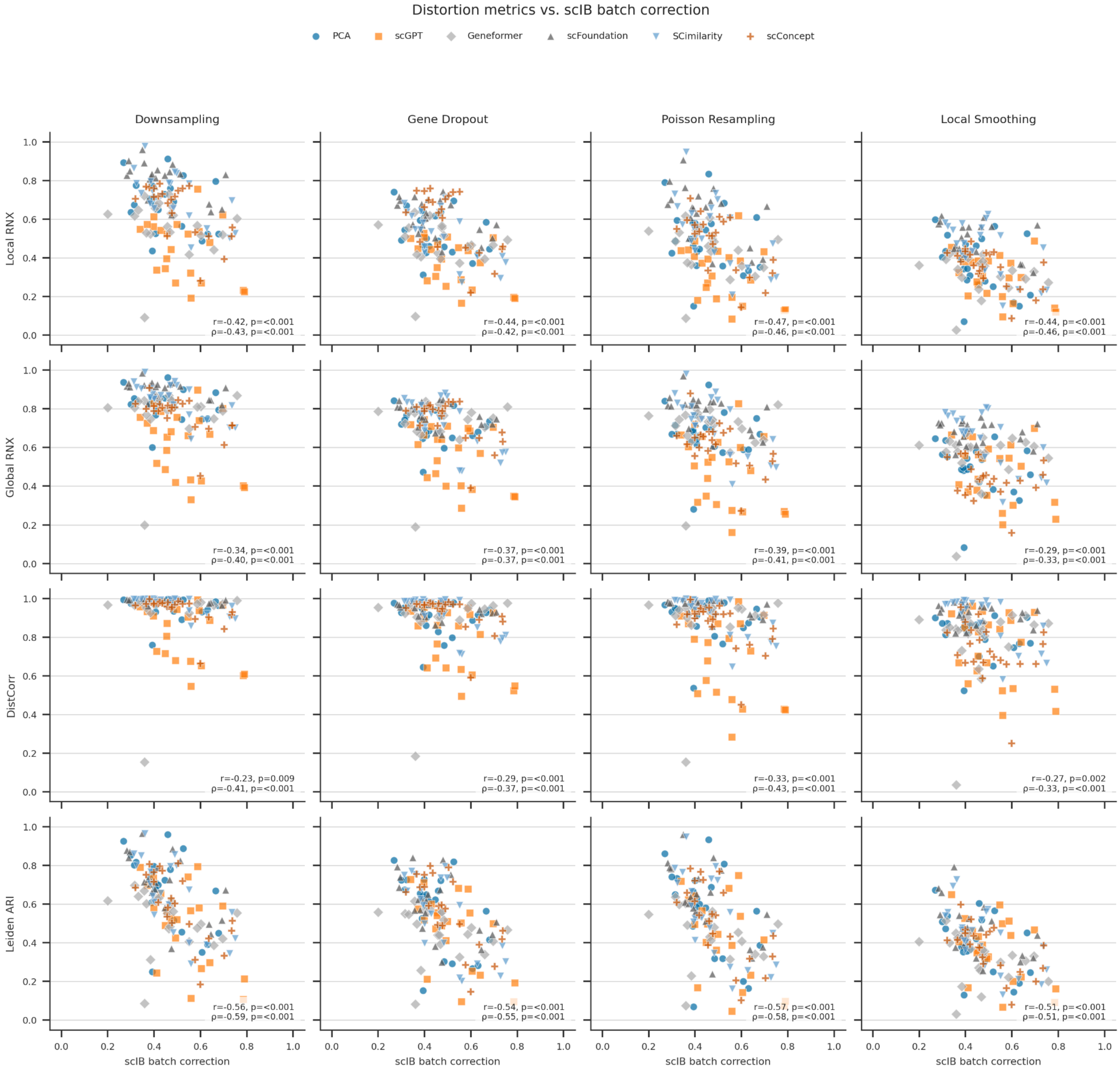


**Figure S13. Label-free distortion metrics vs. scib batch correction score.** Conventions as in Figure S12.

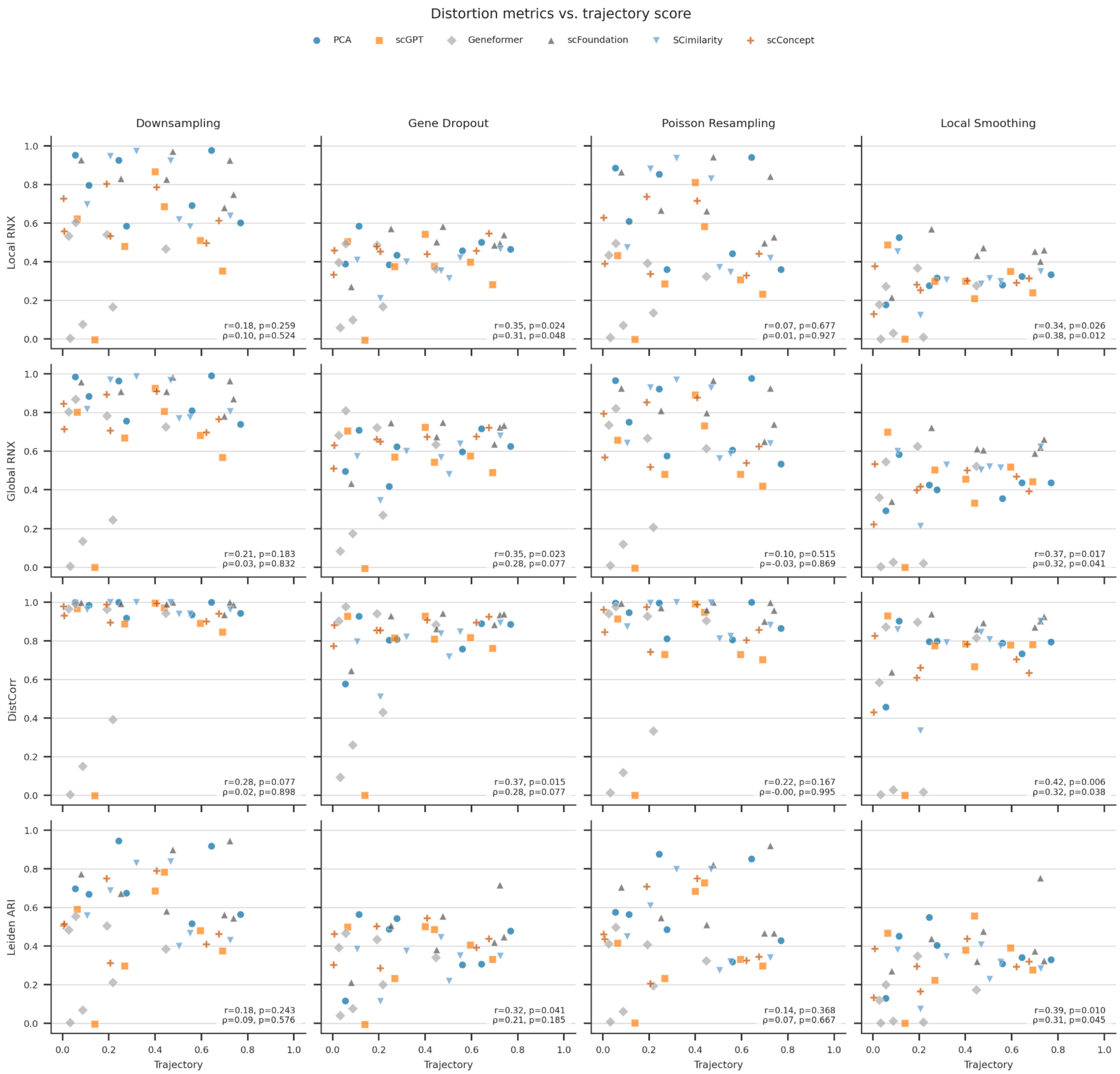


**Figure S14. Label-free distortion metrics vs. trajectory inference score.** Conventions as in Figure S12.

### Supplementary Tables

| perturbation | Free Parameter | Tested values |
|---|---|---|
| Downsample | Fraction | 0.95, 0.9, 0.8, 0.6, 0.4 |
| Poisson resample | Iteration | 1, 2, 3, 4 |
| Gene Dropout | Fraction | 0.1, 0.2, 0.4, 0.6 |
| Local smoothing | K | 5, 10, 15, 25, 50 |

**Table S1.** Perturbations and tested parameter values

| Atlas | Original Size (cells) |
|---|---|
| Arterial | 113,000 |
| Brain | 2,480,956 |
| Immune | 1,821,725 |
| Lung | 2,282,447 |
| Retina | 3,177,310 |
| Tabula Sapiens | 1,136,218 |

**Table S2.** Original atlas sizes used for subsampling validation
